# History of opioid use disorder correlates with altered neuroimmune signatures in the human dorsal root ganglia

**DOI:** 10.64898/2026.10.10.758227

**Authors:** Allison M Barry, Mandee Schaub, Joseph B Lesnak, Erin Vines, Asta Arendt-Tranholm, Úrzula Franco-Enzástiga, Jayden A O’Brien, Ishwarya Sankaranarayanan, Stephanie I Shiers, Arthur Silveira Prudente, Julia R Sondermann, Michael A Wilde, Feng Xian, Tariq Khan, Anna Cervantes, Geoffrey Funk, Peter Horton, Steve Davidson, Temugin Berta, Gregory Dussor, Manuela Schmidt, Theodore J Price

**Author notes:** Corresponding author: Allison Barry (*). co-senior (Manuela Schmidt, Theodore J Price).

## Abstract

Understanding mechanisms for the negative reinforcing effects of opioids are fundamental to improving patient care. Using human dorsal root ganglia (DRG) from donors with and without opioid use disorder (OUD), we set out to examine functional and molecular correlates of chronic opioid use. At a molecular level, we detected disrupted immune signatures throughout cells in the peripheral nervous system (PNS), spanning glial-neuroimmune interactions to changes in non-neuronal senescence. At the physiological level these changes are paired with sensitization of DRG sensory neurons. Our findings reveal a neuro-inflammatory state in the PNS of individuals with OUD. Targeting therapeutics towards these PNS-based mechanisms could be an approach to alleviating the negative reinforcing effects of opioids.

**One Sentence Summary:** Using a combinatory approach of quantitative proteomics, snRNA-seq, and functional readouts in human DRG, we provide evidence of molecular and physiological changes across cell types indicative of neuro-inflammatory signalling in the peripheral nervous system of individuals with opioid use disorder.

## Main Text

As early as the 16^th^ century, there have been written accounts of opioid withdrawal and the pain that accompanies it. In رساله افيون [*Afyunieh*; The Opium Treatise], ابن مسعود شيرازى عماد الدين محمود [Imad al-Din Shirazi] dedicates an entire chapter to opium withdrawal, including a description of “pain of joint and other areas” *(1)*. Centuries later, in one of the most widely read English essays on opioid addiction, Thomas De Quincey wrote that to speak of opium withdrawal “would be indeed *infandum renovare dolorem* [to renew the unspeakable pain]” *(2)*.

These symptoms - now largely referred to as the negative reinforcing effects of opioids - are an important component of opioid dependency and opioid use disorder (OUD). Extensive work has been done in recent decades to better understand the negative reinforcing effects of opioids. There is now a rich literature covering how these symptoms emerge from changes in reward circuitry, as well as abundant evidence for changes in neural pathways related to nociception *(3)*. These involve changes in the immune system, in sensory neurons in the DRG, in microglial signalling in the spinal dorsal horn, and in neuronal circuits related to nociception in the brain *(3–8)*. Many receptor systems have also been associated with these effects, ranging from the canonical opioid receptors (MOR, DOR, and KOR) to toll like receptors (TLRs, mainly TLR4) to other neuromodulatory systems *(9)*. An important gap in this knowledge is how opioid dependency and/or OUD affects the human PNS. This has not yet been studied in detail, and most of what we know about the effects of prolonged opioid administration on the DRG comes from animal studies.

Pain and OUD are closely linked. The majority of individuals with OUD have had or still have chronic pain *(10)*, and opioid use is further implicated in secondary pain states, both from opioid-induced hyperalgesia and from painful withdrawals. In people who inject drugs, 57% of participants cite very or extremely painful opioid withdrawal symptoms, with higher withdrawal pain associated with non-fatal overdoses in the same cohort *(11)*. In parallel, withdrawal-associated injury site pain is a phenomenon where a healed (and typically pain-free) injury site becomes painful during withdrawal. This occurs in almost 20% of all individuals experiencing opioid withdrawal (∼40% of individuals with a previous injury) *(12, 13)*. The descriptions of withdrawal-associated injury site pain are similar to those associated with inflammatory pain, suggesting a mechanism that involves nociceptors in the DRG, and/or neuro-immune interactions with these nociceptors *(14)*.

As we enter the fourth decade of the opioid crisis, morbidity and mortality rates remain high. Opioid overdose remains the leading cause of death in young adults across the USA and Canada, and as of 2022, ∼3.7% of adults in the USA were estimated to have required treatment for OUD *(15–17)*. Central mechanisms of OUD have already been implicated across both sensory and affective components, but aspects of withdrawal pain suggest peripheral involvement *(1, 2, 11–13)*. We set out to explore peripheral mechanisms of OUD. Our goal was to better understand how OUD is associated with molecular and functional changes in the DRG through tissues recovered from organ donors either with or without a history of OUD. Fear of uncontrolled pain is a barrier to deprescription and detoxification *(18)*, so a better understanding of peripheral mechanisms may also inform future pain management in these patients.

## Results

We have previously demonstrated the utility of multi-omic analyses in the human DRG (hDRG) to gain insight into mechanisms of peripheral neuropathy *(19)*. We have now extended this work to donors with OUD to understand the impact of this disorder on the PNS (**Figure 1**). OUD, chronic pain, and alcohol/other substance use status were determined by review of the donor’s medical information using pre-determined set of criteria as noted in **Table 1**. For example, OUD was defined as evidence in these records of: 1) opiate use without pain, 2) prescription opioid use and pain with explicit description of opiate abuse/overdose and/or 3) illicit/street opioid use (eg. heroin) regardless of pain status (**Table 1**). Full cohort metadata are provided in **STable 1**.

**Fig. 1.**
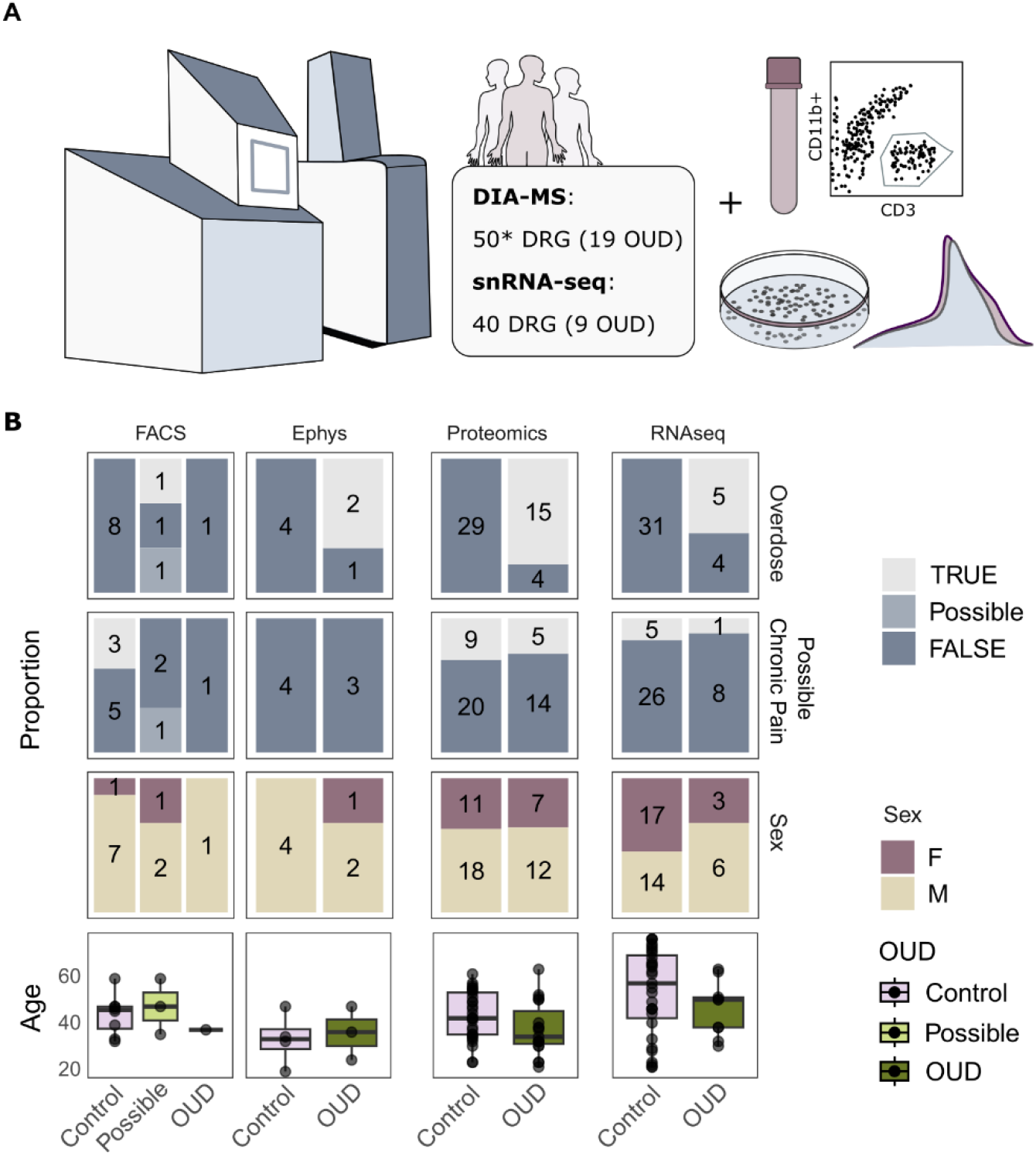
Study Overview. A. Schematic of snRNAseq, DIA-MS proteomics, fluorescence activated cell sorting (FACS), and Electrophysiology (Ephys). B. Cohort details by experiment by proportion with absolute hDRG donor numbers included on each bar. F; female. M; male. TRUE/FALSE; relative to each condition. OUD; opioid use disorder. *2/50 proteomic samples were not considered due to unclear OUD status. Individuals with ‘Possible’ OUD had substance abuse and/or overdose without explicit opioid use, with full cohort definitions Table 1 and metadata in STable 1.

**Table 1.** Cohort Definitions.

|  |  |  |
| --- | --- | --- |
| Opioid Use (OU) | True | explicit use of opioids (any type), regardless of pain/prescription/abuse status |
|  | Unknown/<br>Possible | “pain medication” or “drug/substance use/abuse” where additional description not given |
|  | False | everything else |
| Opioid Use Disorder (OUD) | True | opiate use without pain (any type), or prescription opioid use + pain with explicit description of addiction/opiate abuse/overdose or street opioid (eg. heroin) regardless of pain status. |
|  | Unknown/<br>Possible | “drug/substance use/abuse” where additional description not given or opioid use + pain with hinted but not explicit abuse. |
|  | False | everything else |
| Substance Abuse | True | explicit use of illegal substances (excluding marijuana), or “drug/substance use/abuse” |
|  | Possible | prescription may be abused |
|  | False | everything else |
| Alcohol abuse | True | “daily/heavy alcohol use” or “alcohol/ethanol abuse”, or “beer 6 pack 3-4 times per week” or “binge drinking” |
|  | False | everything else, including “occasional alcohol use”, OR “social drinker” |
| Possible Chronic Pain | True | explicit “chronic [site] pain”, “Arthritis”, any peripheral neuropathy (or diabetes + gabapentin) which may be painful, and/or a description of “back pain” which may or may not be chronic |
|  | False | everything else, including headache disorders |

Human DRG from ∼100 donors were divided across data-independent acquisition mass spectrometry (DIA-MS, 50 donors), single nucleus RNA sequencing (snRNA-seq, 40 donors), as well as *in vitro* functional experiments and flow cytometry with some overlap (**Fig. 1A, STable 1**). Data were age-and sex-matched across molecular datasets, with a subset of individuals having experienced chronic pain in both control and OUD conditions (**Fig. 1B**). Functional experiments were age-matched, with data summarised per donor where relevant due to the lower available number of donors.

### Quantitative proteomics highlights immune disruption in the hDRG

DIA-MS proteomics across 50 donors revealed immune dysregulation in the DRG of individuals with OUD (**Figure 2***, see data availability)*. Our proteomics averaged about 90000 peptides/12500 proteins per sample (**SFig. 1A**), with no clear outliers (**SFig. 1B**) and a consistent coefficient of variation across conditions (**SFig. 1C**). Across 50 donors, 19 had OUD, 5 had opioid use (and pain) without OUD, and 2 had an unclear OUD status (**Fig. 2A**). Broadly, proteomic expression did not show clear separation by condition by PCA (**Fig. 2B**). In line with our previous studies, we obtained good coverage of key ion channels and neuronal markers (eg. *SCN10A*, *TRPV1*, and *PIEZO2)* (**Fig. 2C**) *(20)*.

**Fig. 2.**
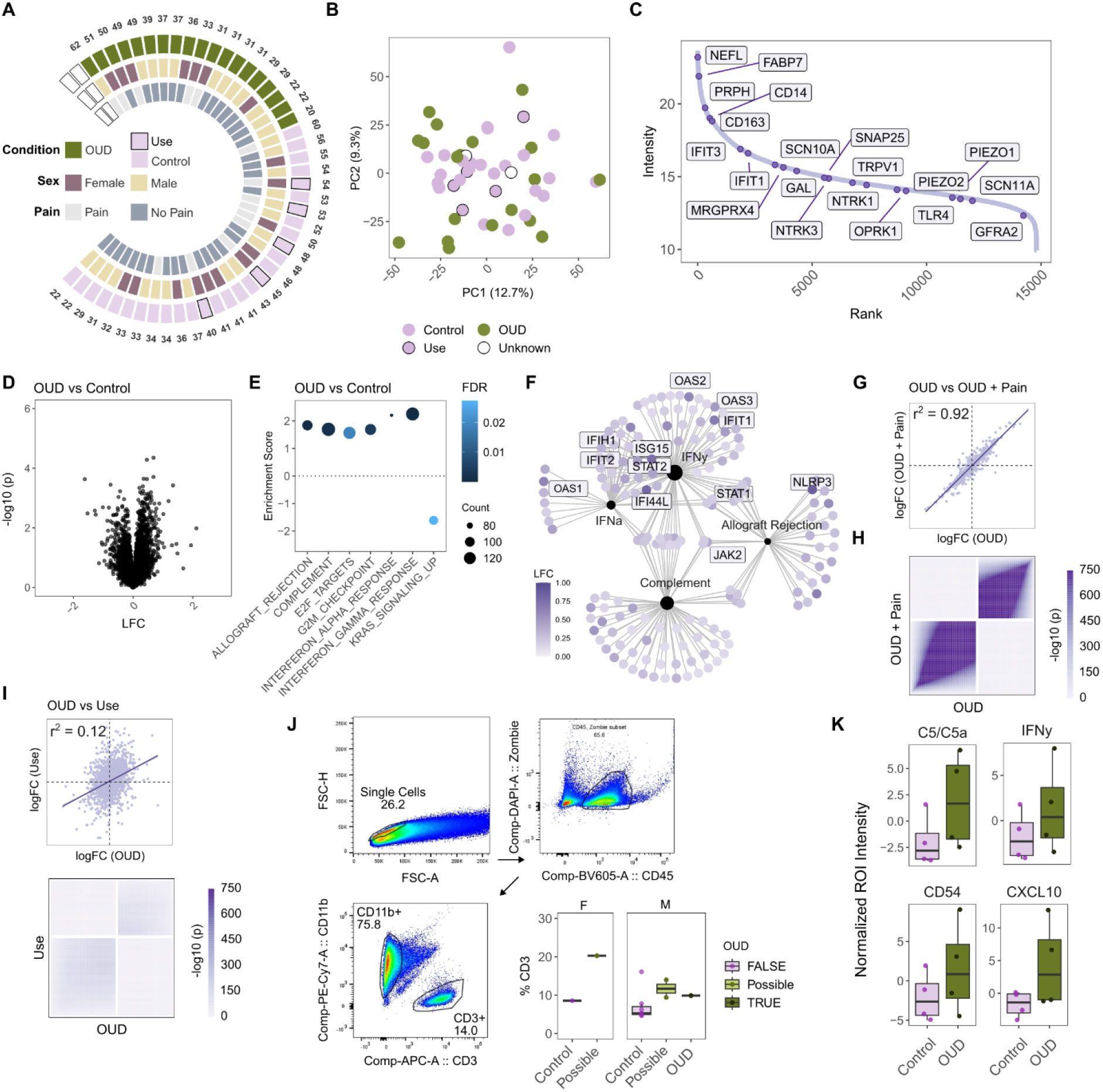
Proteomic signatures of OUD in the hDRG. A. Proteomic cohort metadata, with 2 donors removed due to unknown OUD status (white boxes). B. PCA across 50 DIA-MS hDRG donors. C. Ranked protein expression (median) with key proteins highlighted. D. OUD vs Control, modelled as ∼ Batch + Condition + Sex. E-F. Gene set enrichment analysis (GSEA) for OUD vs Control (E), and gene-concept network plots (F) for significantly regulated pathways, coloured by protein log fold change (LFC). G-H. Sensitivity analyses modelling Pain as a covariate. G. Correlation between fold changes with and without pain modelled. H. Rank-rank hypergeometric overlap (RRHO) test with and without pain modelled. I. Correlation and RRHO test between OUD vs Control and Opioid Use (Use) vs Control. J. Fluorescence activated cell sorting (FACS) for CD3+ T cells in the hDRG (Wilcox p = 0.028 *). K. Cytokine expression across hDRG protein lysates. LFC = log fold change, ROI = region of interest.

While we did not observe differentially expressed proteins in OUD vs Control (∼ Batch + Condition + Sex, **Fig. 2D, STable 2**), we detected an upregulation of immune signalling including hallmark pathways for allograft rejection, interferon alpha- and gamma-response, and complement (**Fig. 2E, STable 3**). Across immune-related pathways, multiple relevant proteins trend towards increased expression, including OAS2/3, NLRP3, and ISG15 (**Fig. 2F**).

A subset of donors with and without OUD had documented pain. To see whether this contributed to the inflammatory signal in OUD, we performed sensitivity analyses by considering ‘pain’ as a factor. Here, we saw high correlation to the original comparison (r^2^ = 0.92) (**Fig. 2G, STable 4**), as well as high concordance via rank-rank hypergeometric overlap testing (**Fig. 2H**). There were no differences in enriched pathways (**SFig. 1D, STable 5**) suggesting that the significant inflammatory signal in OUD is not confounded by pain status.

When looking specifically at donors who used opioids for their clinical care (without OUD), interferon signalling pathways were no long significant (FDR for IFNγ = 0.54, for IFNα = 1), while complement signalling was significantly regulated (**SFig. 1E, STable 6**). Here, modelling ‘pain’ as a covariate abolished previously seen pathway enrichments (∼ Batch + Condition + Sex versus ∼ Batch + Condition + Sex + Pain) (**STable 7**). This suggests we may indeed capture a pain phenotype in this subgroup (opposed to signatures of opioid use without OUD), as all individuals without OUD who used opioids had a concurrent pain phenotype. Comparing non-OUD Use vs Control and OUD vs Control, protein changes were not correlated (**Fig. 2I**). Together, this implies potential differences between the effects of opioids in OUD versus opioid use in a medically controlled, prescribed setting.

We hypothesized that the immune signatures detected in OUD may be due to an increased infiltration of immune cells into the hDRG. Using fluorescent activated cell sorting (FACS), we saw a significant increase in CD3+ T cells in donors with OUD (or possible OUD), compared to age- and sex-matched controls (**Fig. 2J, STable 8,** Wilcox p = 0.028, see Table 1 for cohort definitions). In contrast, CD11b+ cells were unchanged (**SFig. 2**, Wilcox p = 0.279). We also measured corresponding cytokine levels from pooled OUD and control protein samples using a commercial cytokine assay (**Fig. 2K, SFig. 3**). Overall, cytokines were changed across conditions (ANOVA for adjusted_Mean_ROI ∼ Condition*Cytokine, p < 0.001), with no significance in individual cytokines (post hoc Tukey’s HSD for Condition*Cytokine, FDR ∼ 1).

### Transcriptional correlates of OUD

To gain cell-type specific resolution, we turned to snRNA-seq (**Figure 3, SFig. 4***, see data availability)*. From 40 donors (including 9 OUD) (**Fig. 3A, SFig. 4A-B**), nuclei were annotated into 9 main cell types after quality control, with neurons identified by *SNAP25* and *PRPH* expression (**Fig. 3B, SFig. 4C-F**).

**Fig. 3.**
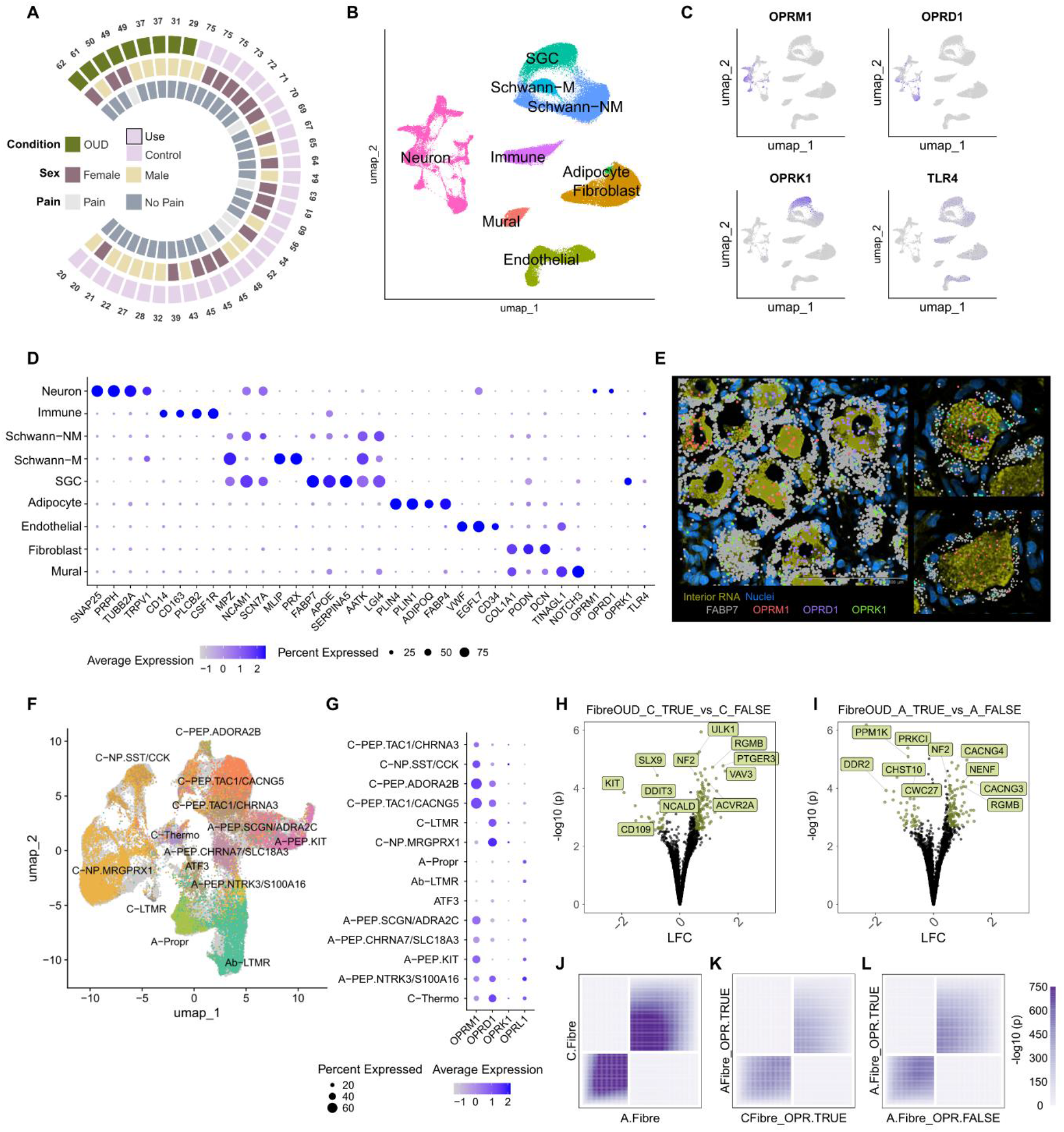
Transcriptional signatures of OUD in the hDRG. A. snRNA-seq cohort metadata. B. UMAP across cell types. C. Opioid receptor expression across cell types. D. Cell type marker expression. E. Xenium transcriptomic validation of opioid receptor expression across neurons and satellite glial cells (FABP7+). F. UMAP across neuronal subclusters. G. Opioid receptor expression across neuronal subpopulations. H-I. Differentially expressed genes (OUD vs Control) for C-(H) and A-fibre (I) populations. J-L. Concordance testing between A- and C-fibre (J), A- and C-fibres expressing opioid receptors (K), and A-fibres with and without opioid receptor expression (L). LFC = log fold change. OPR = opioid receptor.

Molecular correlates of OUD may arise from the direct and/or indirect effect of opioids, thus we first investigated opioid receptor expression across cell types (**Fig. 3C**). *OPRM1* and *OPRD1* show higher neuronal expression than *OPRK1*, with *OPRM1* expression in a subset of human nociceptors as previously described (**Fig. 3C-D**) *(21)*. *OPRK1* was predominantly expressed in human satellite glial cells (SGCs, **Fig. 3C-D**), mirroring reports by Staedtler et al., (2024), with dominant expression in human satellite glial cells (SGCs, **Fig. 3C-D**) *(21)*. Toll-like receptor 4 *(TLR4)*, which has been shown to interact with certain opioids, was also expressed broadly across non-neuronal cells in the hDRG (**Fig. 3C-D**) *(9, 22)*. We validated receptor expression using Xenium spatial transcriptomics, revealing neuronal subtypes with OPRM1 and/or OPRD1/OPRK1, as well as a strong co-localization of FABP7 and OPRK1 in the cells surrounding neurons (**Fig. 3E**).

### Sensory neurons exhibit a distinct molecular signature in OUD

DRG sensory neurons were subset for higher resolution analyses (**Fig. 3F, SFig. 4G**). These barcodes were integrated with the recently published hDRG reference atlas for annotation *(23)*, resulting in 30k neurons across 20 cell populations (**Fig. 3E, SFig. 4H**). Due to the lower neuron count compared to the reference atlas, some populations (eg. C-Thermo subdivisions) were merged into 14 final subtypes with distinct opioid receptor expression profiles (**Fig. 3G**). Differentially expressed gene (DEG) testing was then performed on A- and C-fibre populations, pseudobulked by donor (∼ Condition + Sex) (**Fig. 3H-I, STable 9**) *(24, 25)*. We saw overlapping DEGs across A- and C-fibres, including the BMP co-receptor RGMB (DRAGON), the acid sensing ion channel *ASIC2*, and *IL12A*, which may contribute to IFNγ induction in T cells *(26)*. Other DEGs, like the auxiliary AMPA subunits *CACNG3/4* were only significantly regulated in A-fibres. Broadly, there was high concordance between A- and C-fibre changes in OUD (**Fig. 3J**). This held when comparing only neuronal subtypes expressing *OPRM1* and/or *OPRD1* receptors, with most opioid receptor-expressing sensory populations expressing both in at least 20% of their nuclei (**Fig. 3K**). Furthermore, there was a high degree of similarity between A fibre changes in OUD with and without *OPRM1/OPRD1* receptor expression, suggesting these effects likely do not rely on the activation of these receptors (**Fig. 3L**). Opioid receptors were detected to some extent in all C-fibre populations, thus the effect of direct vs indirect activation through OPRs could not be tested here.

### Non-neuronal cells from human DRG are dysregulated in donors with OUD

To probe non-neuronal changes in OUD we first built an interactome with CellChat across broad cell groupings (**Fig. 4A**). In comparing OUD and control ligand-receptor interaction strengths, we saw an increase in immune signalling towards neurons and SGCs, as well as towards non-myelinating Schwann and mural cells (**Fig. 4B**). In parallel, predicted endothelial and mural signalling towards fibroblasts was increased.

**Fig. 4.**
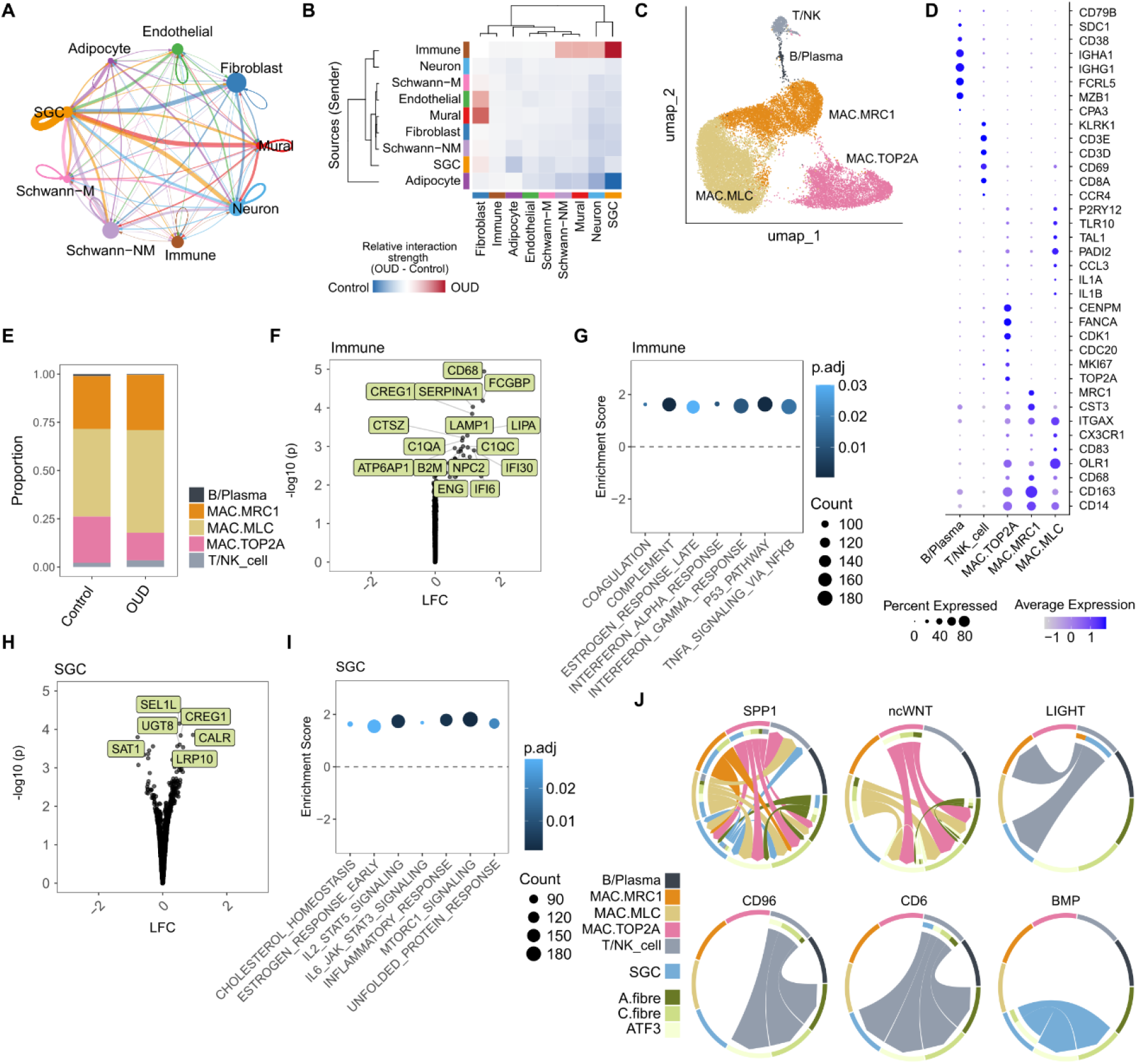
snRNA-seq interactomics in OUD. A. CellChat predicted interactions between cell types, with line widths representing interaction strengths. B. Relative interaction strength change in OUD vs Control. C. UMAP of immune subpopulations. D. Immune subpopulation markers. E. Immune cell proportions across conditions. F. DEGs from pseudobulked immune cell. G. GSEA enrichment within immune cells. H. DEGs in pseudobulked SGCs. I. GSEA enrichment within SGCs. J. Predicted pathway interactions for DEG ligand and/or receptors across neurons, SGCs, and immune subtypes. Arrows indict the direction of communication, from ligand to receptor. p.adj = adjusted p value; LFC = log fold change.

Endothelial and mural cells act as a meningeal neuroimmune interface, can express TLR4, and have roles in chronic inflammation in the CNS *(27)*. Given the initial interactome analyses (**Fig. 4B**), as well as the increase in T cells in donors with OUD (**Fig. 2J**), we next wanted to see if these cells were altered in OUD (**SFig. 5**). We constructed an interactome with DEGs from these subpopulations (**SFig. 5A**), with DEGs from fibroblasts (eg. *ITGB1, COL6A2, AGRN*, **SFig. 5B, Stable 10**), endothelial cells (eg. *DLL4, FRMD4B*, and *CTSB*, **SFig. 5C, STable 11**), and mural cells *(TOGARAM2, SNX10, ACAP1*, **SFig. 5D, STable 12**). Each cell type showed an upregulation of IFNγ, along with various immune-related pathways in fibroblasts and endothelial cells. Angiogenesis was upregulated in fibroblasts (**SFig. 5E-G, STable 13**). The differential expression of various ligands/receptors across these cell types hints at broad changes in meningeal-related cells from donors with OUD (**SFig. 5H, Stable 14**). Many of these interactions hint at basement membrane and extracellular matrix remodelling (COLLAGEN, FN1), others may be involved in leukocyte migration and immune interactions (JAM, CLDN, GRN/AGRN). The upregulation of the receptor *ITGB1* on fibroblasts may affect a variety of pathways, including Collagen, Laminin, JAM, FN1, while the concomitant upregulation of the ligand *COL6A2* as well suggests another role for fibroblasts in the COLLAGEN pathway. Another ligand upregulated on fibroblasts, *SDC2*, may act through *ADGRA3* on mural cells, while the upregulation of *CLDN11* may involve the CLDN pathway within fibroblasts themselves.

### Glial-neuro-immune interactions are predicted to change in OUD

Next, to probe the immune signalling towards neurons and SGCs, we first subclustered our immune nuclei into 5 distinct populations (**Fig. 4C-D**). Three broad macrophage populations were detected, including proliferating macrophages (MAC.TOP2A), microglial-like cells (MAC.MLC), and a general macrophage population (MAC.MRC1) *(23)*. All macrophage populations expressed *TLR4* (**SFig. 6A**). We also identified mixed populations of B cells and Plasma cells (labelled B/Plasma) and T and NK cells (T/NK cells). No significant differences in proportion were detected (**Fig. 4E**, two-way ANOVA, p = 0.8423). We ran DESeq2 by donor across immune barcodes, detecting a significant upregulation of inflammatory genes including *LAMP1, LIPA*, and *IFI6/30* (**Fig. 4F, STable 15**), as well as a corresponding upregulation of inflammatory pathways, including complement signalling as well as IFNα and IFNγ pathways (**Fig. 4G, STable 16**). Using a secondary pathway analysis approach – camera – which further accounts for gene correlations within each pathway (see methods), we could mirror these results, with both type 1 and type 2 interferon signalling upregulated, as well as protein secretion, unfolded protein response, and DNA repair (**SFig. 6B**). Overrepresentation analysis of DEGs highlighted various gene ontology pathways, including lymphocyte mediated immunity, complement activation, IgG binding, and various secretory granule compartments (**SFig. 6C**), indicative of an increased immune response. Within macrophage subpopulations, we detected an increased in protein secretion throughout, with IFNγ and IFNα increases in MAC.MRC1 and MAC.TOP2A populations (**SFig. 6D-F**)

A similar approach was taken for satellite glial cells (SGCs), which may be directly activated by opioids through *TLR4* and/or *OPRK1* (**Fig. 3C-D**). Similar to immune cells, SGCs were transcriptionally different in OUD, with DEGs such as *CALR* and *SEL1L* upregulated (**Fig. 4H, STable 17**). Here, a unique set of pathways were upregulated, including IL2-STAT5 as well as a broad inflammatory response (**Fig. 4I, SFig. 6G, STable 18**). Together, this mirrors the pro-inflammatory state detected with bulk proteomics, while suggesting unique OUD signatures by cell type in the human DRG.

To test how these individual cell type changes may affect one another through differentially expressed ligands and receptors, a higher resolution interactome was built with CellChat (**Fig. 4J, STable 19**). Some pathways likely involve many cell types, while some are predicted to have more specific pairings. For example, the ligand SPP1 is upregulated in immune cells, and may interact across immune, neuron, and SGC subtypes. The receptor *LTBR* is also upregulated on immune cells, with predicted input from T/NK cell ligands in the LIGHT pathway *(TNFSF14* to *LTBR)*.

Neuronal DEGs may also have implications for glial-neuro-immune interactions: First, *FZD3* was specifically upregulated in C-fibres, which may receive input from *WNT5A*, *WNT5B* and/or *WNT11* expressed on macrophages. Second, the receptor *ACVR2A* was upregulated on both C- and A-fibres in donors with OUD, with predicted input from SGCs through various BMP signalling ligands. Third, the upregulation of *ALCAM* and *NECTIN1* on A-fibres may be relevant for T/NK cell signalling through CD6 and CD96 pathways respectively.

### Non-neuronal cells show evidence of senescence

There is evidence OUD can accelerate aging in the nervous system *(28–30)*. To test whether any cells from donors with OUD exhibit signs of senescence, we ran GSEA against the ‘Sen Mayò gene set, comparing OUD vs Control *(31)*. A subset of non-neuronal cells showed positive enrichment for senescence in OUD, including SGCs, endothelial cells, fibroblasts, and proliferating macrophages (MAC.TOP2A) (**SFig. 7**).

### Kappa opioid receptors exhibits a unique expression profile in human DRG

Kappa signalling in the CNS has previously been linked to opioid withdrawal *(8)*. Even so, *OPRK1* expression shows clear species differences: OPRK1 is predominantly expressed in rodent sensory neurons while in humans, higher expression is seen in SGCs with more sparse neuronal expression (**Fig. 3C-E**) *(21, 32)*. Staedtler et al. (2024) have previously described non-neuronal *OPRK1* expression in human SGCs using both commercial and custom *in situ* probes *(21)*. Here, we have replicated their human SGC expression using snRNA-seq and Xenium spatial transcriptomics (**Fig. 3C-E**). To verify the species difference, we further examined OPRK1 expression in a recently published cross-species atlas (**SFig. 8**) *(33)*. In line with our snRNA-seq and Xenium data, *OPRK1* was detected in human SGCs (**SFig. 8B**). Minimal expression of *Oprk1* was found in non-neuronal DRG cells across mouse, rat, guinea pig, cynomolgus macaque, and rhesus macaque (**SFig. 8C**). This mirrors RNAscope staining, with *OPRK1* expression in human SGCs across six additional donors (3M + 3F, **SFig. 8D-E, STable 20**), and *Oprk1* expressed in a subset of murine neurons but not SGCs (**SFig. 8F**).

### Sensory neurons from donors with OUD are hyperexcitable

We hypothesized that DRG neurons recovered from donors with a history of OUD might be hyperexcitable due to changes in glial-neuroimmune signalling *(14, 34, 35)*. To test this, we performed patch clamp electrophysiology in age-matched primary hDRG cultures from donors with and without OUD (**Figure 5**). DRG neurons recovered from donors with a history of OUD (n= 3 OUD, 4 Control donors) showed similar median capacitance, but a trending reduced median rheobase in OUD donors (Wilcox, p ∼ 0.2) (**Fig. 5A-C**). While the firing threshold in OUD neurons was similar to controls (**Fig. 5D**) OUD neurons generated significantly more action potentials when stimulated with a depolarizing ramp protocol (Wilcox, p < 0.05), suggesting a sensitized state (**Fig. 5E**). On the other hand, we did not observe differences in the rate of spontaneous activity (**Fig. 5F**), nor in other action potential properties (rise and decay slops, half width, or peak amplitude) across neurons (**Fig. 5G-K**, n = 27 OUD, 75 Control neurons). These experiments suggest that sensory neurons from donors with OUD may be sensitized but are not spontaneously active (as seen in many patients with neuropathic pain) *(36, 37)*.

**Fig. 5.**
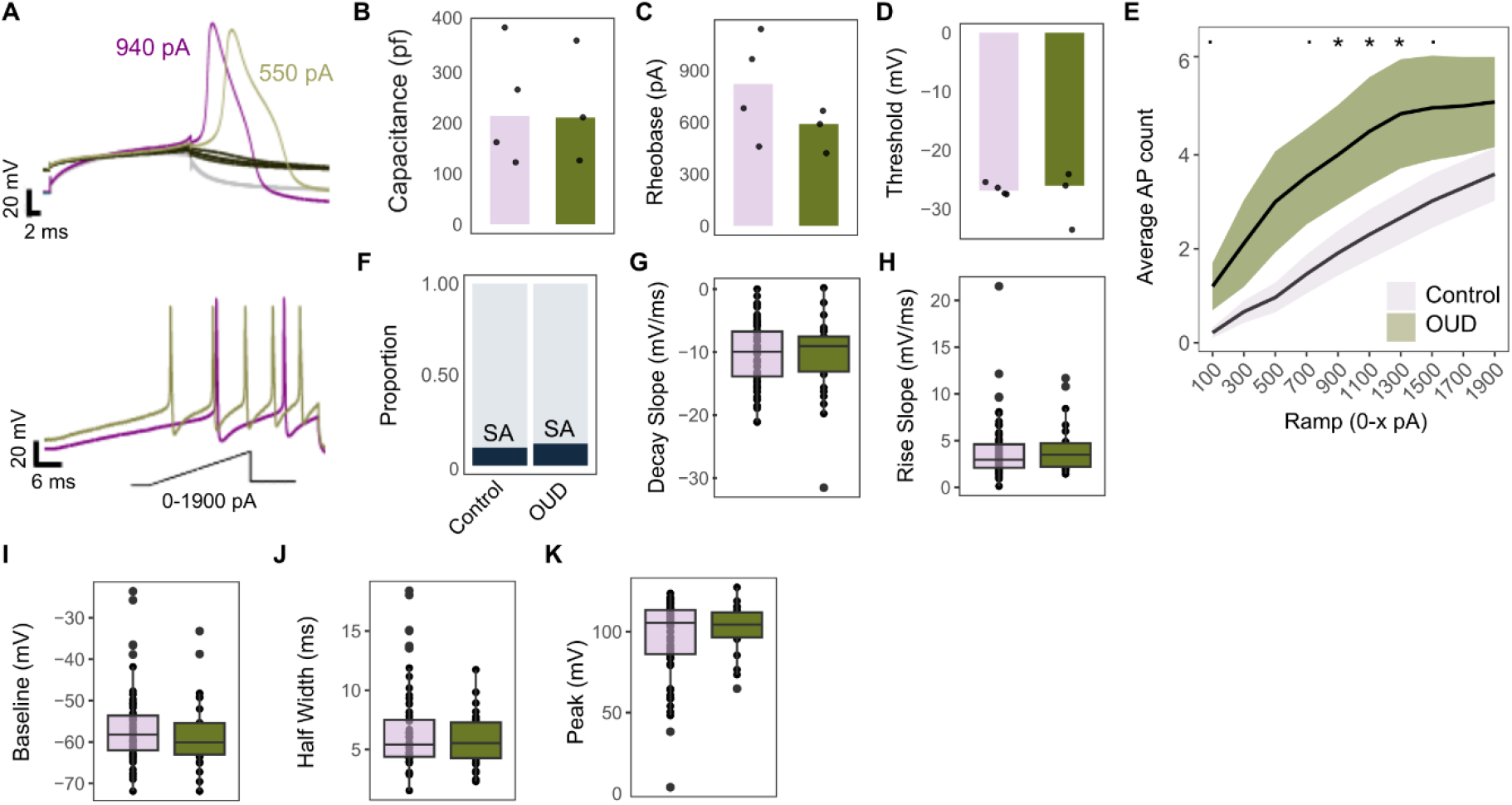
Functional characteristics of hDRG sensory neurons in donors with and without OUD. A-K: Baseline electrophysiology characteristics, L-O: Calcium Imaging. A. Representative action potential traces (top), and representative traces to the ramp protocol, from 0-100 pA to 0-1900 pA (bottom). B. Median capacitance by donor, C. Median rheobase, D. Median threshold potential (mV). E. Mean action potential (AP) response to a ramp protocol, with standard error highlighted. F. Total spontaneously active (SA) neurons per condition. G-K. Action potential properties by neuron. G. Decay Slope. H. Rise Slope. I. Baseline mV. J Action potential half width (ms). K. Peak amplitude (mV). Wilcox, * p < 0.05,. p < 0.1. Donors; n= 3 OUD, 4 Control. Neurons; n = 27 OUD, 75 Control.

## Discussion

In the early 19^th^ century, the English essayist and critic Thomas De Quincey referred to his opium withdrawal as “internal rheumatism”, hinting at an inflammatory pain comprising the rheumatic definitions of the time *(2, 38)*. To our knowledge, our study is one of the first to directly assess the possibility that in humans, OUD induces a peripheral inflammatory state affecting the DRG. We find that DRGs recovered from people with OUD exhibited features of an inflammatory state across omics, with neurons with and without opioid receptor expression exhibiting similar transcriptional changes indicative of indirect effects. Neurons from these individuals also showed signs of sensitization. This neuronal activity is consistent with signatures of inflammatory pain, such as that associated with rheumatism, and distinct from the neuronal signatures observed in neuropathic pain, which include marked spontaneous activity in a subset of neurons *(36, 37, 39)*.

Though correlative, our findings may provide a neurophysiological substrate associated with the negative reinforcing effects of opioids, originating through peripheral sensory systems. Broad cytokine increases and immune disruption have previously been reported in plasma and CNS in individuals with OUD *(5, 40, 41)*, and we extend this here to the PNS. We hypothesize that non-neuronal opioid signalling through *TLR4*, as well as immune cell and immune mediator infiltration, may be driving these effects: *TLR4* is widely expressed in non-neuronal cells in the DRG, and is a known, non-canonical opioid receptor with a role in CNS neuroinflammation *(9, 22)*. Opioid-based activation of glial *TLR4* triggers a pro-inflammatory response, which includes microglial and astrocyte activation paired to cytokine release and altered glutamate homeostasis respectively *(9, 22, 42–44)*. While less studied in the context of opioids, *TLR4* activation on endothelial cells can further alter blood brain barrier permeability, again contributing to neuroimmune signalling through changes in vascular permeability *(9, 45)*.

In the periphery, DRG neurons can be directly sensitized during chronic opioid exposure via MOR *(46, 47)*, and *TLR4* activation by low-dose morphine can sensitize both human and rodent sensory neurons *(48, 49)*. Neuroimmune signalling in the DRG has also been discussed preclinically in the context of chronic opioid exposure, albeit to a lesser extent than in the CNS. CGRP and Substance P levels increase in cultured rat primary afferents after morphine exposure *(50)*, and opioid induced hyperalgesia has been linked to chemokine signalling via SGCs in Sprague-Dawley rats *(51, 52)*. Both MOR and *TLR4* blockers can attenuate the development of opioid induced hyperalgesia in rats *in vivo*, suggesting relevance for both MOR and *TLR4* pathways *(52)*.

We detect largely concordant changes in sensory neurons with and without opioid receptor expression and do not detect changes in opioid receptor expression with OUD, thus we hypothesize neuronal changes to be an indirect consequence of local, chronic inflammation. DRG neurons from donors with OUD showed an increased rate of action potential firing compared to age-matched controls, with no changes in spontaneous activity. This mirrors neuronal activity in inflammatory pain states, including that seen in response to rheumatoid arthritis and/or in response to inflammatory mediators *(34, 53)*. Even so, our study is underpowered to firmly interpret these functional experiments. To achieve a power of 0.8 (α = 0.05) for rheobase alone, we would require 11 donors per group. Even with overdose as a leading cause of death in young adults *(15, 16)*, it would take years to reach this sample size given the history and medical records from organ donors we can recover tissues from through our pipeline. Given the urgency in addressing the opioid crisis now, we feel these trends are worth reporting. Excitability was predominately increased in sensory neurons from donors with OUD once neurons had started firing action potentials, indicating that the mechanisms controlling repetitive firing in are likely altered in the OUD population we sampled from. Extrapolating clinically, this would suggest that these individuals may have reported hyperalgesia, but not pain at rest, a phenotype that is linked to chronic inflammatory pain *(14, 54, 55)*. These findings complement a recent meta-analysis on patients receiving opioid agonist treatment for OUD, where patients showed hypersensitivity to cold pain *(56)*.

Non-neuronal cells were also broadly affected in donors with OUD. Across molecular dimensions, we measured increased complement signalling, as well as increased type I and II interferon signalling in the DRG. The significant increase in CD3+ T cells in OUD vs Control DRG may directly relate to the increased interferon signatures, but more work is needed to untangle which T cell subpopulations may be driving this. IFNγ-secreting regulatory T cells (Tregs) have previously been described in blood samples from patients with OUD, with preclinical evidence these can infiltrate the nervous system with chronic opioid exposure *(5, 41)*. Individuals on maintenance therapy for OUD have also shown restored plasma Treg levels, suggesting at least some of these effects may be reversible in patient populations with current treatment options *(41)*.

We anticipate parallels between the human PNS and CNS, as bulk RNA-seq of post-mortem brain tissue from individuals with OUD showed a similar pattern of neuroinflammation and interferon signalling to the DRG *(40)*. People with OUD have also been shown to have significantly reduced blood–brain barrier integrity versus controls *(29)*, with white matter changes suggesting reactive/activated glia *(9, 22).* Preclinically, opioid withdrawal has also been linked to blood brain barrier permeability changes: in mice taking opioids, this permeability change allows fragile-like Treg infiltration and IFNγ secretion in the nucleus accumbens, stimulating synaptic remodelling and contributing to withdrawal behaviours *(5)*.

The increased senescence in non-neuronal PNS cells further fits into the narrative of an ‘aged’ nervous system in OUD: In the CNS, this includes evidence of Alzheimer’s-like plaque formation *(28, 29)* and cognitive decline *(57)*. In (non-OUD) Alzheimer’s disease models, interferon signalling, complement, and vascular senescence have been linked to amyloid β plaque formation *(58, 59)*. Due to the relative ease of accessing peripheral over central nervous tissue in humans, DRG dysfunction and senescence in OUD may further provide a window into related CNS disturbances in people with OUD.

### Limitations and future directions

Human DRG tissue is difficult to access, and while we have made great strides to increase the amount of tissue and data available, we are still limited in our experimental capacity in this area. For this study, cohort definitions are based on available medical histories (see Methods). Given the complex nature of OUD and polysubstance abuse disorders, as well as significant limitations in health care access among this population *(17, 60)*, we cannot discount that this history is incomplete, eg. for an underlying pain condition or confounding substance, which may contribute to changes in neuronal excitability and/or immune profiling. We also note that hDRGs are taken from donors on life support, where tissues may be in a withdrawal process. Even so, results published from preclinical model systems and human plasma suggest that immune dysfunction in OUD is not limited to individuals on life support *(5, 41, 42, 61)*.

Moving forward, we invite researchers to develop an accurate, humanized model system that can test causal effects of OUD. Longitudinal, large-scale monitoring of individuals with OUD for developing pain and immune-centric conditions will also help address the clinical consequences of immune disruption. If plasma signatures are found to correlate to changes at the level of the DRG, this would provide an additional layer of cohort stratification from more deeply phenotyped participants.

## Conclusion

The negative reinforcing effects of opioids have long been studied in the context of the CNS, or, in the case of the PNS, in preclinical model systems. Here, we provide evidence for an “inflammatory pro-nociceptive-like” state in the human DRG in individuals with OUD. This includes molecular changes at both the transcriptomic and proteomic levels, as well as function sensitization at the level of the neurons. By providing molecular correlates of OUD that relate to nociception, as well as broad support for neuroimmune changes in the nervous system of individuals with OUD, we demonstrate clear, molecular and cellular evidence for PNS dysfunction in people with OUD

## Methods

### Human tissue procurement

University of Texas at Dallas: All procedures were approved by the University of Texas at Dallas Institutional Review Board (protocol Legacy-MR-15-237). The Southwest Transplant Alliance obtained informed consent for research use through first-person authorization (e.g., driver’s license or legal documentation) or from next of kin. Donor screening and consent policies are governed by the United Network for Organ Sharing, and the Alliance adheres to CDC standards and undergoes biannual inspection by the Department of Health and Human Services. Medical history was summarized by the Southwest Transplant Alliance, obtained from next of kin and/or recent hospital records as available. All donor information was handled in compliance with HIPAA regulations.

Lumbar and thoracic hDRGs were obtained from human organ donors in collaboration with the Southwest Transplant Alliance (Dallas, TX). DRG were surgically recovered within 4 hours of cross-clamp, as previously described *(62)*. Cohorts were defined using criteria outlined in **Table 1**. Due to donor availability, the time course of OUD was not considered, with 3 OUD donors having “prior” OUD listed (1 snRNA-seq, 1 proteomics, 1 FACS; one of these donors had tested positive for opioids on admission). No donors were characterized as opioid dependent without addiction, and no donors were positive for human immunodeficiency virus (HIV-1).

Immediately following recovery in the operating room, hDRGs were buried directly in crushed dry ice for rapid freezing (proteomics and transcriptomics) or in media (for functional downstream experiments). For functional experiments, hDRGs were placed either in cold artificial cerebrospinal fluid for less than 1 hour or in calcium-free BrainBits Hibernate A medium supplemented with N2 Supplement-A (1%), NeuroCult SM1 (2%), penicillin/streptomycin (1%), GlutaMAX (1%), 2 mM sodium pyruvate, and 0.1% bovine serum albumin, and stored at 4 °C for 4–16 hours. We have previously demonstrated the effects of Hibernate A medium have minimal effect *(63)*.

University of Cincinnati: The collection of dorsal root ganglia (DRG) from deidentified organ donors was reviewed by the Institutional Review Board of the University of Cincinnati (#00003152, Study ID 2015–5302) and was deemed to be exempt. DRG (L4, L5) were collected in the operating room within 90 min of aortic cross clamp and after removal of vital organs. DRG were placed in N-Methyl-D-glucamine-artificial cerebrospinal fluid (aCSF) solution at 4°C and were dissected as described *(64)*.

For RNAScope (below), DRG were immersion fixed in 4% paraformaldehyde in phosphate-buffered saline (PBS) overnight at 4°C, cryoprotected in 30% sucrose for 24 h at 4°C, and embedded in Optimal Cutting Temperature (OCT) compound (Tissue Tek). Frozen sections (14 µm sections) were mounted onto Superfrost Plus slides (Fisher), dried (15 min) and stored at −20°C.

Donor metadata is provided in Supplemental Table 1.

### Murine colony management

Adult CD1 mice (Stock No: 22, males and females, 8-12 weeks) were purchased from Charles River Laboratories. Mice were housed four per cage at 22 ± 0.5°C under a controlled 14/10 h light/dark cycle, with food and water available *ad libitum*. All experimental procedures were approved by the Institutional Animal Care and Use Committee at the University of Cincinnati (#25-01-27-01), in accordance with the National Institute of Health Guide for the Care and Use of Laboratory Animals. All results are reported according to Animal Research: reporting of in vivo Experiments (ARRIVE) guidelines *(65)*. Animals were assigned randomly to different experimental groups. Sample sizes were determined based on our previous similar studies *(66)*. Investigators were blind to animal treatments, no adverse effects were observed during these studies, and all animals were included in statistical analyses.

### Proteomics

The proteomics data used here is also analysed in parallel by Sankaranarayanan et al, stratified for diabetic peripheral neuropathy *(19)*. Briefly, data-independent acquisition mass spectrometry (DIA-MS) was performed on hDRGs from 50 donors with mixed medical histories as previously described *(67)*, with 2 samples omitted from comparisons due to unknown OUD status (n = 48, with 19 OUD).

Samples were processed in 2 batches, with 8 donors in batch 1 (previously published, *(20)*) and an additional 42 donors in batch 2 representing a diverse group of individuals with OU and OUD, as well as mixed pain and neuropathy conditions, controls across adulthood.

Spectral deconvolution was performed across all samples using DIA-NN (1.9.1, match-between runs enabled). MaxLFQ was calculated using the diann package (R) as previously described *(68, 69)* and the technical replicates for batch 1 were averaged by donor to generate a protein expression matrix across donors. Data were log_2_ transformed prior to downstream analyses.

The batch effect was removed from the protein expression matrix prior to plotting using ‘limma::removeBatchEffect’ *(70)*. PCA was performed on proteins expressed in all samples.

Differentially expressed proteins were tested using ‘limmà. Proteins were considered significant with an FDR < 0.1 and a log2 fold change > 0.5. Pathway analyses were subsequently performed with ‘limma::camerà modelling for OUD + Sex, as well as ‘clusterProfiler::GSEÀ*(70–72)*. Sensitivity analyses were performed for Sex and Age, as well as a broad, possible chronic pain grouping.

### Cytokine Assay

Cytokine levels were measured using the remaining hDRG protein samples, pooled into biological replicates (4 donors per replicate (16 per condition), n = 4 OUD, 4 Control). Instructions for the commercial kit, Proteome Profiler Human Cytokine Array Kit (R&D Systems, Catalog # ARY005B), were followed with slight modification: each membrane was washed independently using 7 ml Wash Buffer in 60×15 mm culture dishes. Membranes were imaged with a ChemiDoc MP Imaging Platform (Bio Rad).

Image quantification was performed in ImageJ. Regions of Interest (ROIs) of equal size were processed using the ROI Manager, and quantification was considered as x̅ _(pixel intensity, PBS)_ - x̅ _(pixel intensity, ROI)_ for each cytokine site, using the PBS negative control per membrane as the background.

### Transcriptomics, snRNA-seq

For nuclei isolation, frozen hDRG tissue was minced into ∼1 mm fragments in ice-cold isolation buffer (0.25 M sucrose, 150 mM KCl, 5 mM MgCl₂, 1 M Tris pH 8.0, 0.1 mM DTT, protease inhibitor, 0.1% Triton X-100, and 0.2 U/µl RNase inhibitor). The sample was then gently homogenized on ice using a Dounce homogenizer (∼15 strokes). The homogenate was passed through a 100 µm strainer and centrifuged at 500 ×g for 10 minutes at 4 °C. The resulting nuclei pellet was resuspended in 2 ml of buffer (1% BSA in PBS, 0.2 U/µl RNase inhibitor) and centrifuged again at 500 ×g for 5 minutes at 4 °C. The pellet was immediately fixed using the 10x Genomics Chromium Fixed RNA Profiling kit for 16 hours at 4 °C, followed by labelling with the 10x Fixed RNA Feature Barcode kit for 16 hours. Library preparation was subsequently performed according to the manufacturer’s protocol.

Libraries were sequenced in multiple batches on either the Illumina NextSeq 2000 at the University of Texas at Dallas Genome Core or the NovaSeq X at Psomagen. Sequencing reads were processed and aligned to the human reference genome (GRCh38) using Cell Ranger v7 (10x Genomics). Ambient RNA removal was performed with SoupX, and doublets were removed using scDblFinder *(73, 74)*.

snRNA-seq data were then processed with Seurat V5 using default parameters unless otherwise described. Data were filtered for fewer than 5% mitochondrial reads, normalized across all samples (’NormalizeDatà) before scaling (’ScaleDatà), integrating with CCA or Harmony (’IntegrateLayers(method = CCAIntegration, orig = “pca”)’. Initial clustering was performed with a resolution = 1, with additional doublet clusters removed manually based on the co-expression of different cell type markers. Data were then reprocessed (from normalization) with the remaining barcodes, clustering at a resolution of 0.5 for cell type annotations.

### Differential expression testing, transcriptomics

DESEq2 was used to test differentially expressed genes (DEGs) by Cell Group and/or subpopulation after pseudobulking by Donor.ID with ‘Seurat::AggregateExpression’*(24)*. The number of barcodes per donor (per cell type) and count matrix were extracted. While DESeq2 can account for library size differences in bulk RNA-seq datasets, the wide range of barcodes per donor limits it’s use in snRNA-seq datasets of this size. Data were thus filtered to remove donors with low barcode counts (neurons> 250 barcodes; neuronal subpopulations/Immune/SGC/Fibroblasts/Mural/Endothelial > 125 barcodes) before modelling ∼OUD + Sex. Genes were filtered for those with fewer than 100 counts, and fold changes were moderated by ‘apeglm’*(24, 25)*. Genes were considered significant with an FDR < 0.1 and an abs(LFC) > 0.5.

### Pathway analyses

Pathway analyses were performed in three ways: 1. For comparisons with DEGs (as defined above), ‘enrichGÒ was used to run an overrepresentation analysis of DEGs against a background of all genes tested in that comparison *(72)*. Redundant Gene Ontology (GO) pathways were removed using ‘clusterProfiler::simplify(cutoff=0.7, by=“p.adjust”, select_fun=min, measure = ‘Wang’)’and pathways were considered significant at an FDR < 0.05. 2. Gene set enrichment analyses (GSEA, via clusterProfiler::GSEA) was performed on ranked fold changes (after apeglm moderation), against the collection of Hallmark pathways, with an FDR < 0.05 considered significant *(72)*. 3. To account for inter-gene correlations among –pathways, ‘limma::camerà was used as a complement to GSEA *(71)*. Pseudobulked count data were filtered for genes expressed in 50% of samples, and were modelled by condition (∼ OUD). Again, an FDR < 0.05 was considered significant.

### CellChat

An exploratory CellChat analysis was run using default parameters across all cell groups for OUD-TRUE and OUD-FALSE independently *(75)*. Interaction strengths were visualized and clustered by weight to determine downstream comparisons. More specific analyses were then run between relevant sender-receiver groups (eg. Immune, Neuron, SGC) after annotating subpopulations. Again, objects were processed for OUD-TRUE and OUD-FALSE independently prior to downstream comparison. Here, ‘ identifyOverExpressedGenes ‘ was not used, due to its inflated false positives when considered each barcode a sample *(76)*. ‘ [[“features”]] ‘ and ‘[[“features.info”]]’ were manually provided as the pseudobulked, DESeq2 output per cell group. In each case, pct.1 and pct.1 were set to 1. A and C fibre comparisons were directly provided, whereas ATF3 had too few barcodes for an independent analysis. DEGs were thus set as a general, pseudobulked neuron comparison.

To explore ligand-receptor interactions where either L/R were considered DEG, ‘ variable.both = FALSE ‘ was set for ‘identifyOverExpressedInteractions()’. Communication was also filtered for ‘min.cells = 100’.

### Rank-rank hypergeometric overlap

Rank-rank hypergeometric overlap (RRHO) was run using the RRHO2 library in R against both limma and DESeq2 outputs using ranked ‘-log10(p-values) × sign(effect size)’, in line with published recommendations *(77)*.

### Xenium Transcriptomics

Xenium spatial transcriptomics was performed on fresh-frozen lumbar DRG tissue obtained from 1 control donor according to 10X published protocol. Tissue was OCT embedded and cryosectioned (10 µm) onto a Xenium slide. Sections were fixed and permeabilized with 4% PFA and 70% methanol according to published 10X User Guide CG000581 Rev D.

Slides were processed with the Xenium in Situ Gene Expression workflow with Cell Segmentation Staining (10x Genomics, User Guide CG000749) with a custom 472-gene panel optimized for DRG. The protocol included overnight probe hybridization, ligation of circularizable probes, enzymatic amplification of gene-specific barcodes, staining to facilitate cell segmentation (ATP1A1/CD45/E-Cadherin for cell boundary, 18s for interior RNA, alphaSMA/Vimentin for interior protein) followed by autofluorescence quenching and staining for DAPI. Imaging was performed on the Xenium Analyzer and transcripts were subsequently visualized and exported via Xenium Explorer (v4.0.0).

### DRG culturing

Human DRGs were dissociated as previously described *(63)*. Briefly, excess connective tissue, fat, and nerve roots were removed to expose the ganglion bulb. Ganglia were then mechanically and chemically dissociated in 1 mg/mL Stemxyme I, 0.1 mg/mL DNase I, and 10 ng/mL recombinant human β-NGF in calcium- and magnesium-free HBSS at 37°C. The suspension was strained (100 μm) and layered (10% BSA in HBSS) before resuspension in prewarmed DRG culture medium consisting of BrainPhys medium supplemented with penicillin/streptomycin (1%), NeuroCult SM1 (2%), GlutaMAX (1%), N2 Supplement-A (1%), 10 ng/mL β-NGF, 2% fetal bovine serum, and 0.1% of a solution containing 5-fluoro-2′-deoxyuridine (3 mg/mL) and uridine (7 mg/mL).

### Electrophysiology

To explore the functional implications of OUD on human sensory neurons, we curated electrophysiological properties across OUD and non-OUD controls from available in house recordings *(63)*. From 25 donors with available electrophysiological data and corresponding medical histories, 12 had vehicle and/or bath recordings available (11M, 1F). From this, a stratified cohort was built blinded to recording data, excluding three individuals with alcohol and/or polysubstance abuse without explicit OUD, as well as two young donors to produce an age-matched comparison (n = 3 OUD (9-15 cells), 4 Control (11-27 cells)). As the only female donor had OUD, it was not possible to produce a sex-matched cohort.

DRG neurons were plated at ∼50 cells per well on 12 mm coverslips in 24-well plates. On the day of recording, cells were transferred to bath solution (135 mM NaCl, 10 mM glucose, 10 mM HEPES, 5 mM KCl, 2 mM CaCl₂, 1 mM MgCl₂, 20 mM sucrose; pH 7.4, 310–320 mOsm) and recorded within 90 minutes. Whole-cell patch clamp recordings were performed using a Multiclamp 700B amplifier with pClamp10 software. Glass pipettes (1–3 MΩ) were filled with intracellular solution (135 mM KCl, 10 mM HEPES, 4 mM Mg-ATP, 0.9 mM Na-GTP, 0.5 mM EGTA, 5 mM NaCl; pH 7.4, 305–320 mOsm). After achieving a gigaohm seal and whole-cell configuration, capacitance was compensated.

Cell health was first assessed in voltage clamp using a −70/0/−70 mV step protocol. In current clamp, spontaneous activity was recorded for 90 seconds. Rheobase was determined using 20 ms current steps (Δ10 pA). Excitability was further assessed using 1 s ramp currents (0–100 pA, increasing by 200 pA up to 2100 pA).

Data were analysed manually using Clampfit 10.4. Excitability and action potential properties are plotted as median value per donor. For the ramp protocol, action potential counts were averaged across all cells per condition and plotted as mean + SEM.

### Fluorescence Activated Cell Sorting

In line with electrophysiology, FACS data were collected in line with Lesnak et al., (2025) *(63)*. Age- and sex-matched controls were used based on the available OUD data. Three donors with unclear OUD status (Overdose and/or drug history without explicit OUD) were included as “Possible” due to the low n, see **Table 1**. In brief, hDRGs were dissociated using enzymatic digestion (Stemxyme I, DNase I, β-NGF), filtered, and centrifuged. Red blood cells were lysed, and myelin was removed using magnetic beads and column separation. Cells were then stained with a live/dead dye, Fc receptor blocker, and antibodies against CD45, CD11b, and CD3. Following washes, cells were sorted by FACS to isolate live CD45 immune populations, including CD11b and CD3⁺ subsets.

### *In situ* hybridization (i.e. RNAscope)

Mice were deeply anaesthetized with isoflurane and transcardially perfused with PBS followed by 4% paraformaldehyde (PFA). Lumbar DRG tissues were isolated and post-fixed in PFA for 1-2 hours before incubation overnight in a 30% sucrose solution. Tissues were then embedded in OCT medium (Tissue-Tek) and cryosectioned at a thickness of 14 µm.

Adult human DRGs were blocked and fixed overnight at 4°C in 4% PFA in PBS (pH 7.4) and incubated in 30% sucrose overnight at 4°C. Tissue was embedded in Tissue Plus (Fisher Healthcare), cryosectioned at 14 μm, and stored at −80°C.

RNAscope was performed following the manufacturer’s instructions using the RNAscope Multiplex Fluorescent Reagent Kit v2 (Advanced Cell Diagnostics, Cat. No: 323110). Probe against the mouse Oprk1 (Cat. No: 316222) and human OPRK1 mRNA (Cat. No: 556711) were purchased from Advanced Cell Diagnostics. For some experiments, RNAscope signals were combined with immunofluorescence, which was carried out as described below.

The cryosectioned tissue sections were washed with PBS followed by incubation in blocking solution (2% of BSA in PBS) for 1 hour and then incubated overnight at 4°C with an anti-HepaCAM (Ab, Novus Biologicals #MAB4108, 1:2,000). Sections were then incubated for 1 hour at room temperature with anti-mouse Alexa Fluor 594 (1:1,000; Thermo Fisher Scientific). Images were acquired using the Keyence BZ-X800., with 2 images per donor (3F, 3M) quantified.

## Data Availability

The mass spectrometry proteomics data have been deposited to the ProteomeXchange Consortium via the PRIDE partner repository with the dataset identifier PXD071905 *(78)*. Raw and processed proteomic data will be publicly available on the SPARC Portal (RRID:SCR_017041). Transcriptomic data will be made available at dbGaP on publication. Proteomic data are searchable under the ‘Proteomics’ tab on https://sensoryomics.com.

## Code Availability

Code will be available at https://github.com/aliibarry/hDRG-oud.

## List of Supplementary Materials

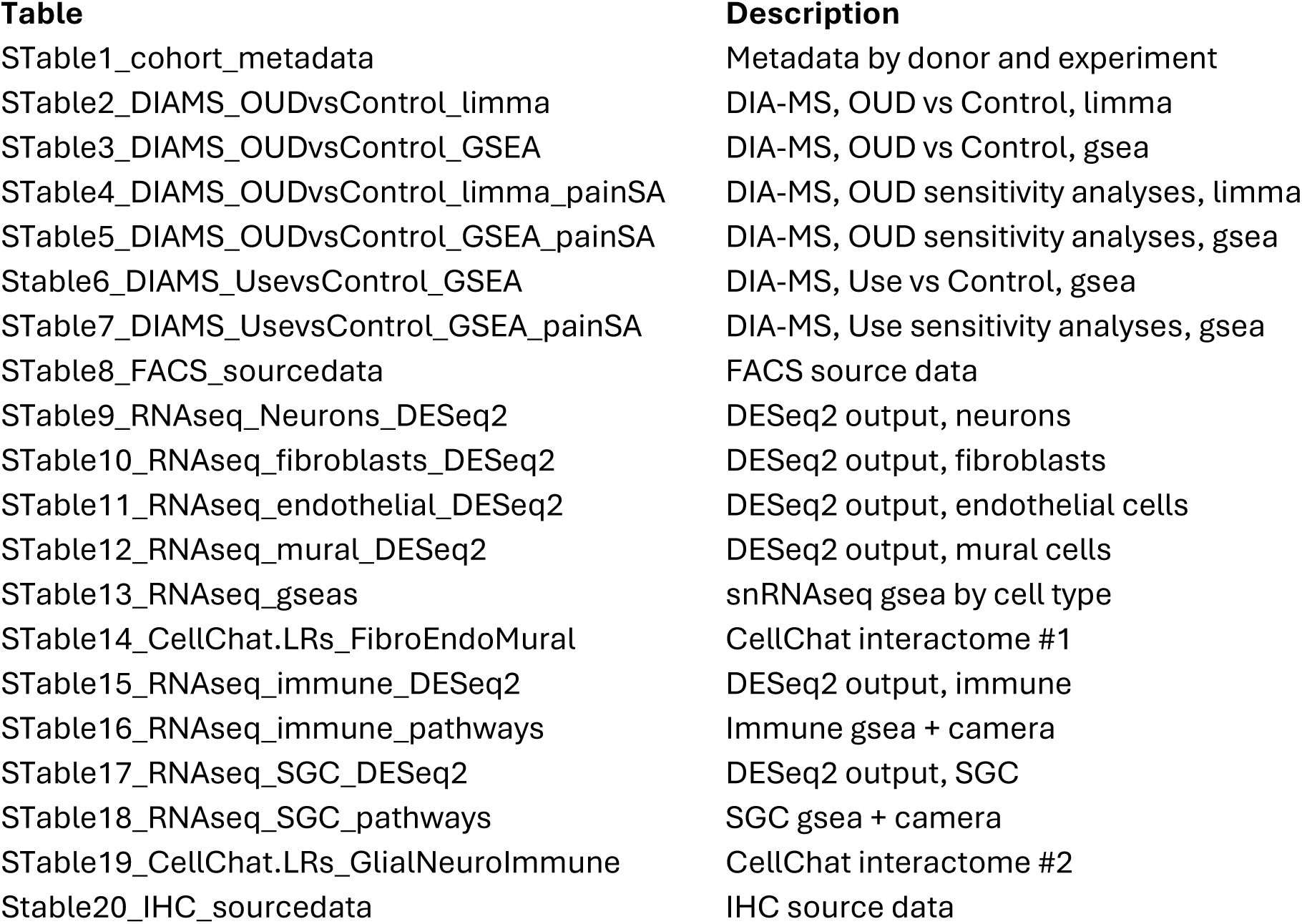

## Abbreviations

CCA: canonical correlation analysis
CNS: central nervous system
DEG: differentially expressed gene
DEP: differentially expressed protein
DIA-MS: data independent acquisition mass spectrometry
DIV: days *in vitro*
DRG: dorsal root ganglia (hDRG – human DRG)
FACS: fluorescence activated cell sorting
FDR: false discovery rate
GSEA: gene set enrichment analysis
HBSS: hank’s balanced salt solution
LFC: log_2_ fold change
NGF: nerve growth factor
NK: natural killer cell
OU: opioid use
OUD: opioid use disorder
PFA: paraformaldehyde
PCA: principal component analysis
PNS: peripheral nervous system
SGC: satellite glial cell

## Author contributions

Conceptualization: AMB, MS, TJP

Methodology: AMB, MS, JBL, AAT, UFE, IS, SS, ASP, JRS, FX, EV, TK, AC, GF, PH, SD

Investigation: AMB, MS, JBL, AAT, IS, ASP

Visualization: AMB, IS, ASP

Funding acquisition: TB, GD, MS, TJP

Project administration: TJP

Supervision: TB, GD, MS, TJP

Writing – original draft: AMB, TJP

Writing – review & editing: all authors

## Acknowledgements

The authors acknowledge the organ donors and their families for their generous gift of life. They thank the Southwest Transplant Alliance, and the members of the Price laboratory for their assistance with DRG recovery. The authors are grateful to the University Vienna – Bruker Center of Excellence for Metaproteomics for their support with DIA-MS. Computation was achieved in part with support from High Performance Computing at The University of Texas at Dallas (HPC@UTD), as well as from the Vienna Scientific Cluster (VSC).

The work was supported by the National Institute of Neurological Disorders and Stroke through the NIH PRECISION Human Pain Network under the award U19NS130608-01 (TJP) as well as through RF1NS136108 (TB). This project was also funded in part by the University of Vienna and the Austrian Science Fund 10.55776/PAT2329024, 10.55776/PAT3085725 and 10.55776/P36554 (MS).

## Generative AI

No generative AI was used in the presentation or writing of this manuscript.

## Supplemental Materials

**SFig. 1.**
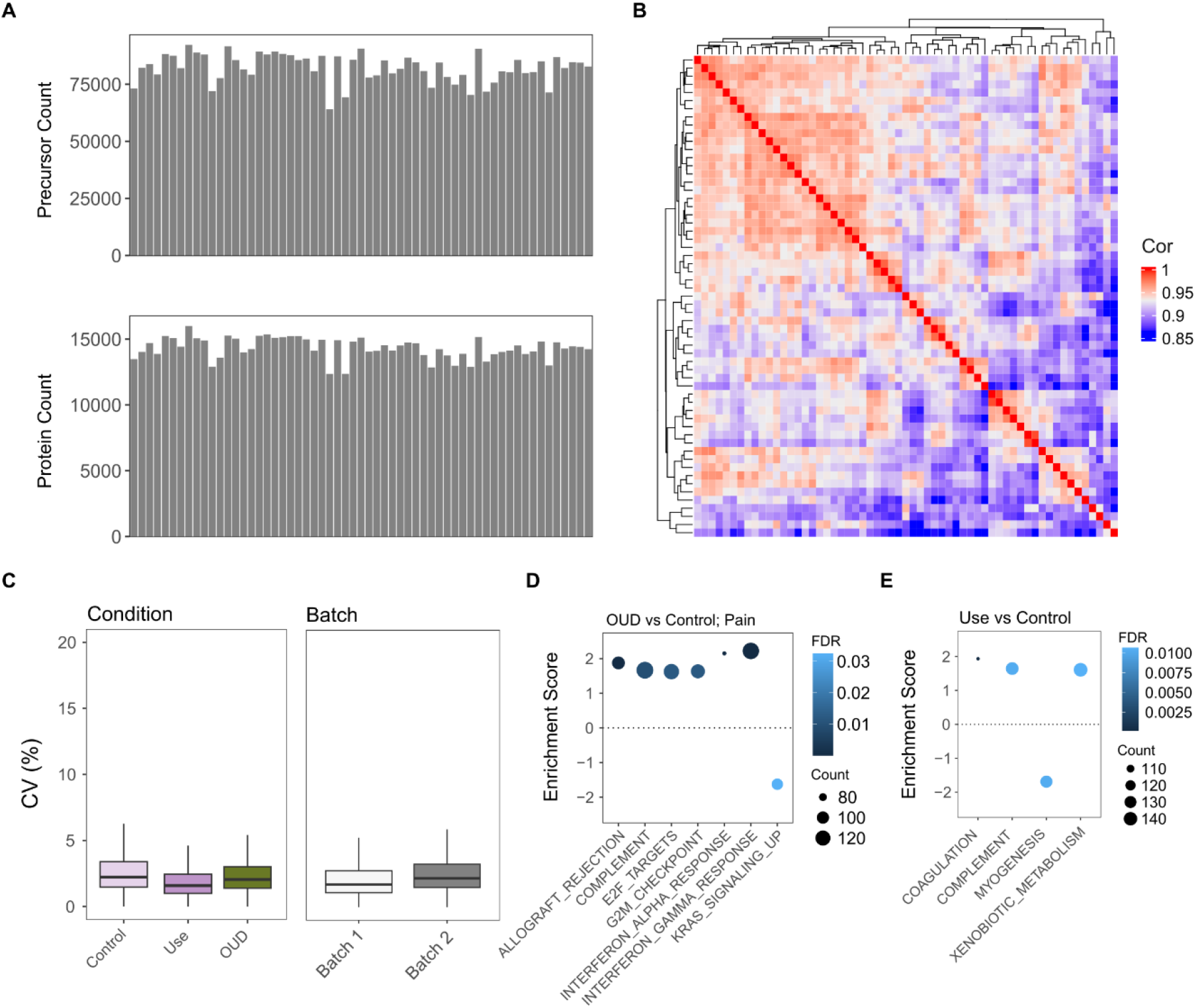
Human DRG proteomics supplement. A. Precursor (top) and Protein (bottom) count per sample. B. Correlation across samples. C. Coefficient of variation (CV) by condition and batch. D. Gene set enrichment analyses as a sensitivity analyses for OUD vs Control, modelling ∼ Batch + Sex + Condition + Pain. E. Gene set enrichment analysis for Opioid Use vs Control (∼ Batch + Sex + Condition). No pathways were significant when Pain was also modelled.

**SFig. 2.**
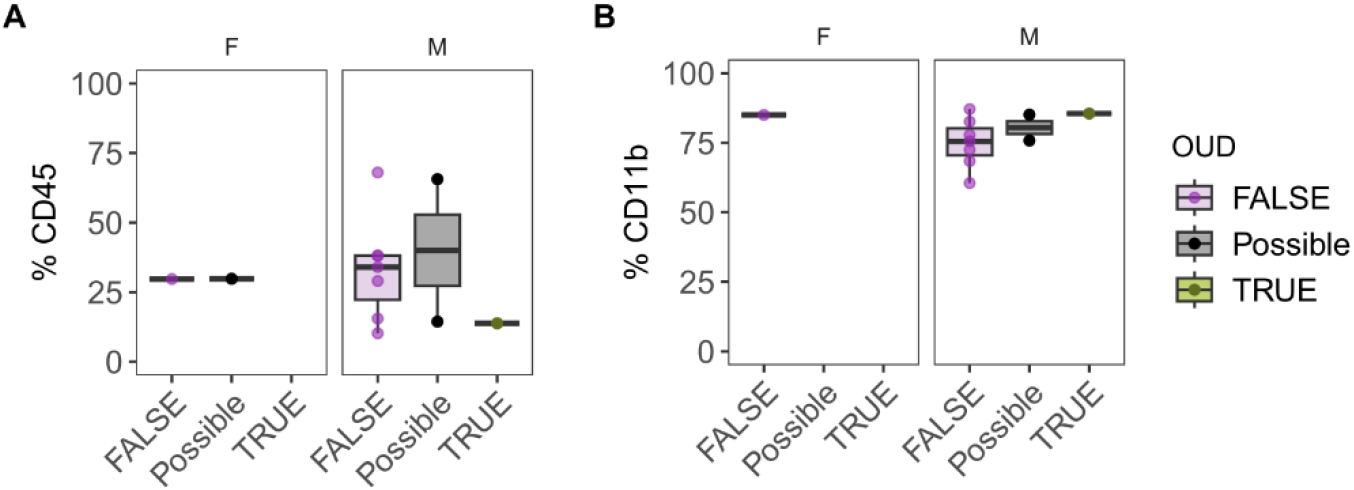
Fluorescence-activated cell sorting (FACS) for immune cells in the hDRG. A. Percentage of CD45 cells per donors. B. % of CD11b + cells. Supplement to Figure 2.

**SFig. 3.**
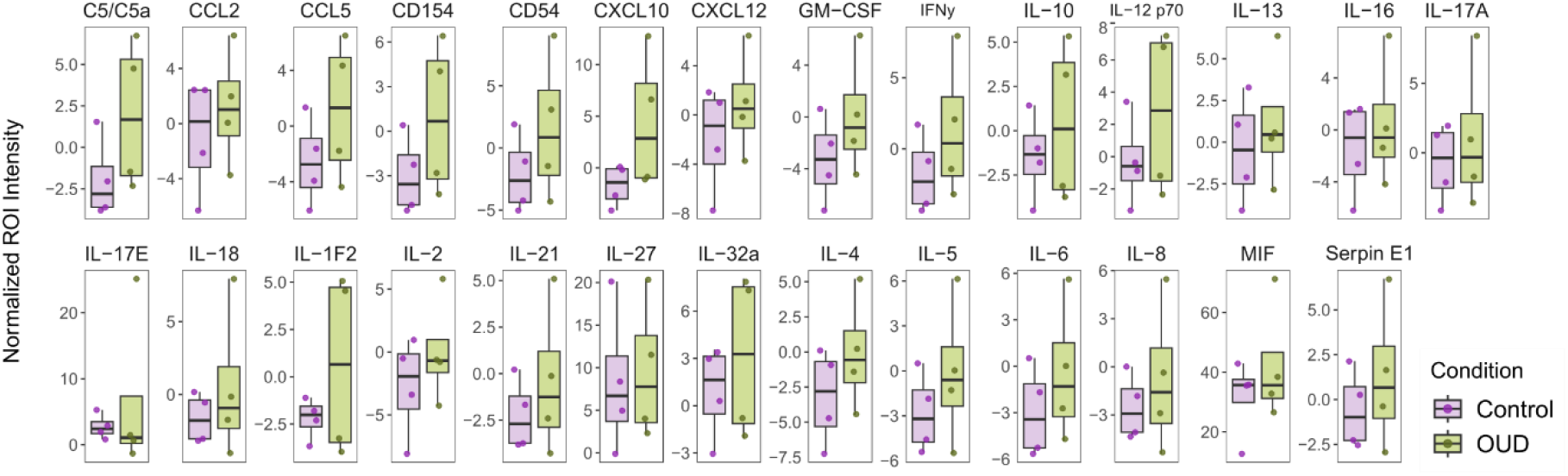
Cytokine expression in hDRG from donors with and without OUD. Cytokine expression across samples (n = 4 OUD, 4 Control, pooled from 4 unique donors each). Supplement to Figure 2.

**SFig. 4.**
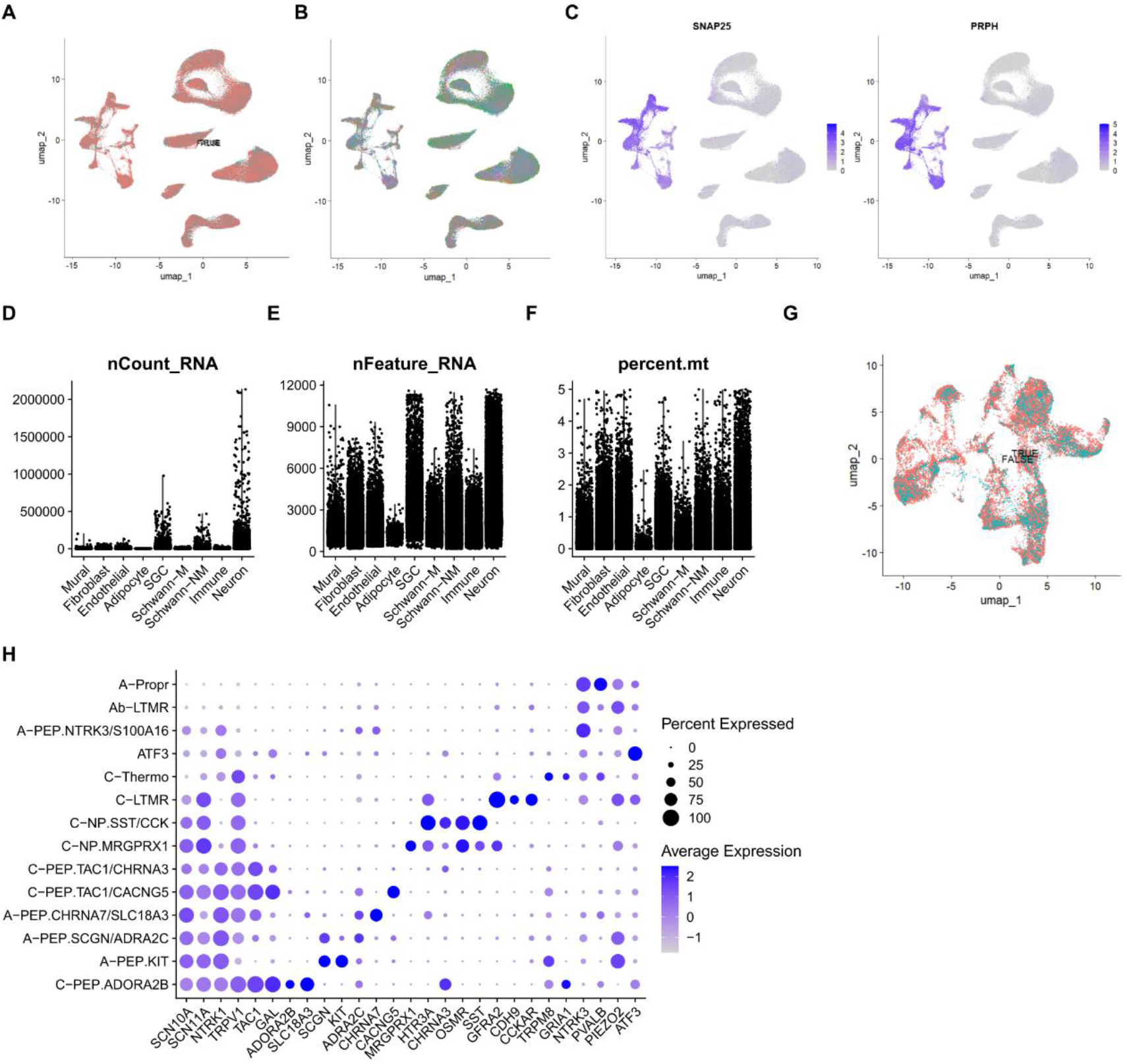
snRNA-seq quality control. A. UMAP by condition. B. UMAP by donor. C. *SNAP25* and *PRPH* expression in the neuronal cluster. D. Read count by cell type. E. Gene count by cell type. F. Percentage of mitochondrial genes. G. Neuronal UMAP by condition (OUD TRUE/FALSE). H. Neuronal subtype markers.

**SFig. 5.**
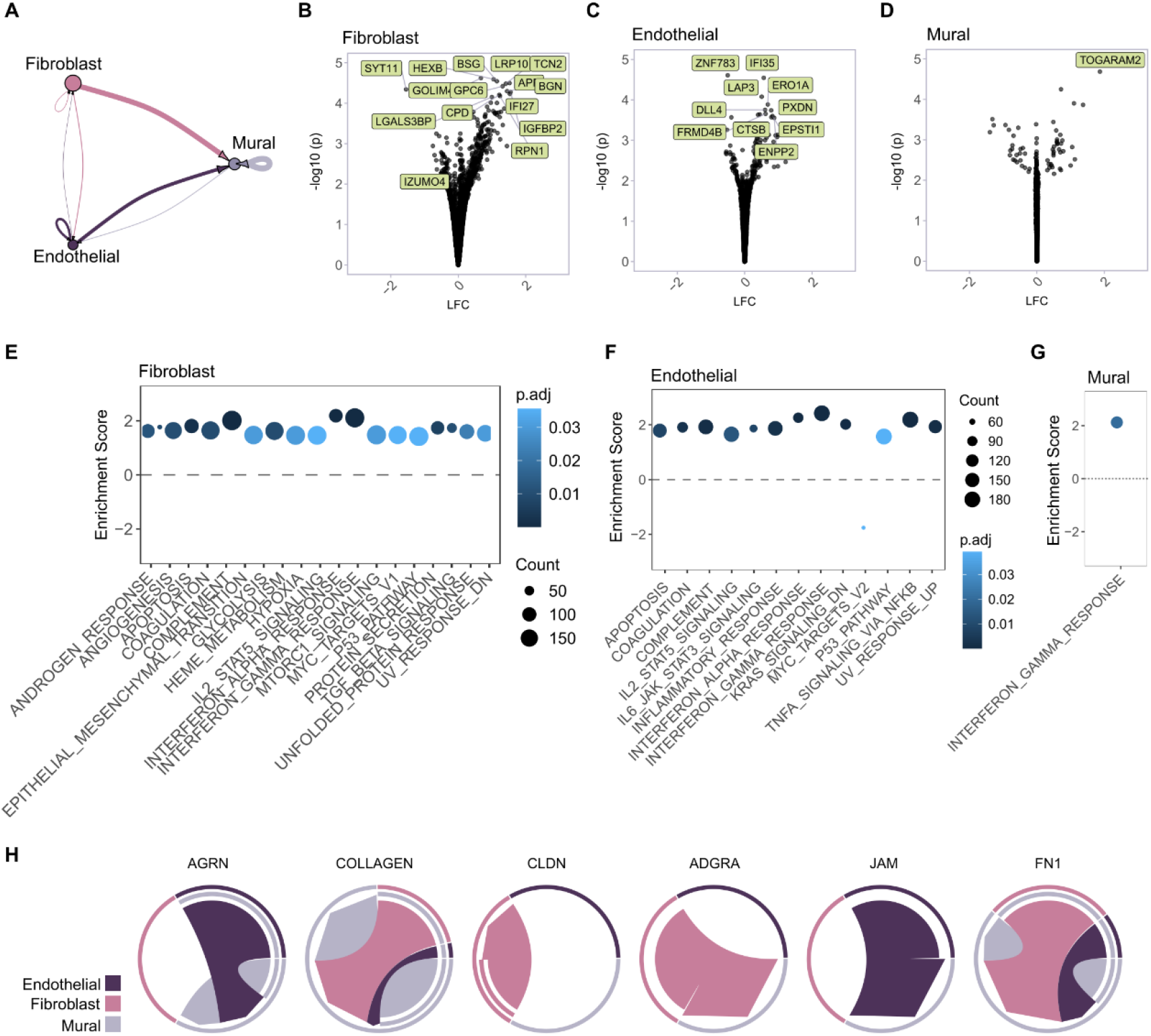
OUD correlates to changes in vascular cells. A. CellChat interactome between fibroblasts, endothelial cells, and mural cells with line weight representing relative weight of interactions. B-D. Differential gene expression (DESeq2 + apeglm on pseudobulked count data by donor) for fibroblasts (B), endothelial cells (C) and mural cells (D). E-G Gene set enrichment analyses for fibroblasts (E), endothelial cells (F) and mural cells (G). H. Predicted interaction pathways for DEG ligands and/or receptors.

**SFig. 6.**
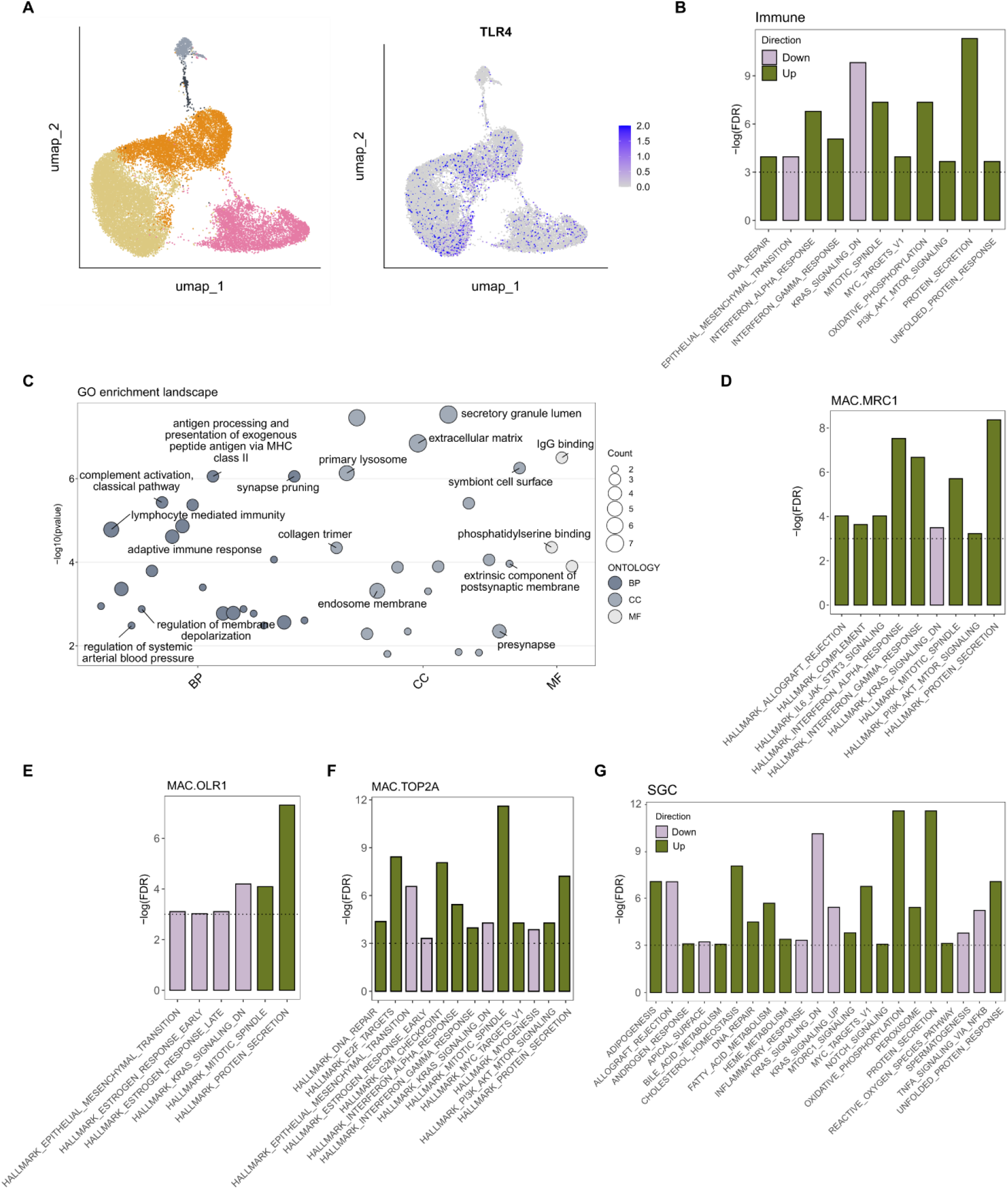
Extended pathway analyses for SGCs and Immune cells. A. Immune UMAP with TLR4 expressed by barcode. B. Immune CAMERA against Hallmark pathways. C. Immune overrepresentation analyses against all 3 gene ontology categories (BP, biological pathways; CC, cellular component; MF, molecular function). D-F. Immune subpopulation CAMERA. G. SGC CAMERA against Hallmark pathways.

**SFig. 7.**
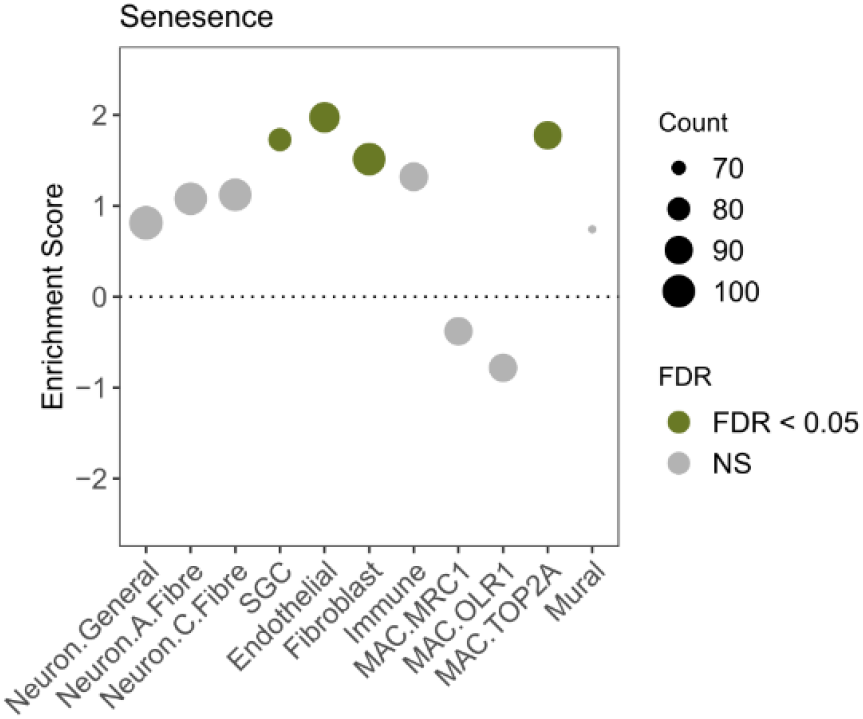
Senescence in OUD. GSEA against the Sen Mayo senescence gene set (OUD vs Control), by cell type.

**SFig. 8.**
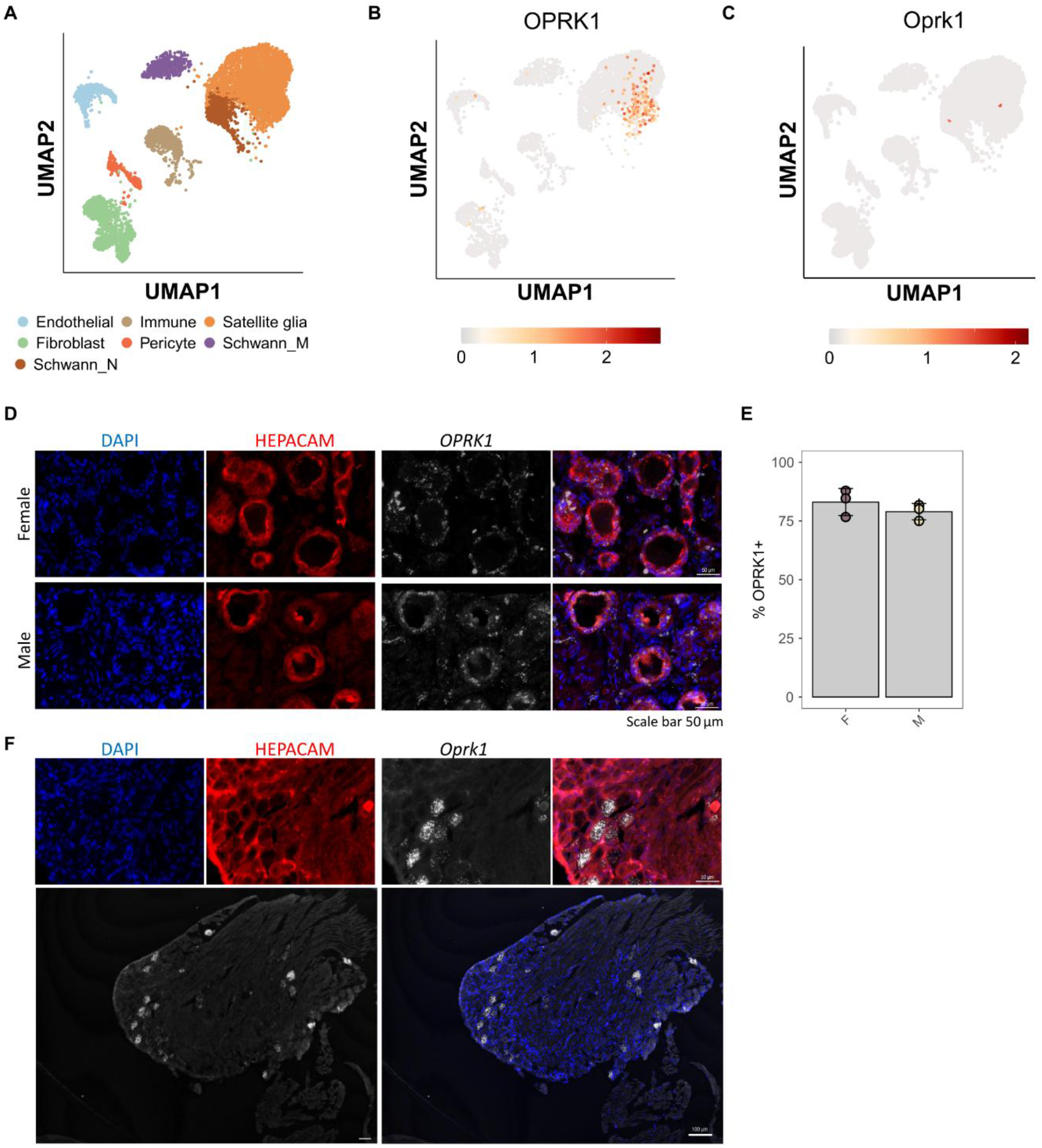
*OPRK1* expression in human satellite glial cells. A. Integrated, cross-species single cell and single nuclear RNA-seq for non-neuronal cells in the DRG. B. *OPRK1* expression in human non-neuronal cells. C. *Oprk1* expression across non-human primates and rodents. D. RNAscope for *OPRK1* and the SGC marker HEPACAM in human DRG (3F, 3M). E. *OPRK1* quantification of HEPACAM+ cells (% *OPRK1* positive). F. RNAscope for *Oprk1* and the SGC markers HEPACAM in mouse DRG. 10x scale *Oprk1*, 50 µm. 10x scale merged, 100 µm. 40x scale, 50 µm.

## References

1. A. Moosavyzadeh, F. Ghaffari, S. H. Mosavat, A. Zargaran, A. Mokri, S. Faghihzadeh, M. Naseri, The medieval Persian manuscript of Afyunieh: the first individual treatise on the opium and addiction in historyJ. Integr. Med. 16 (2018), doi:10.1016/j.joim.2018.02.004.

2. T. de Quincey, Confessions of an English Opium-Eater (Taylor and Hessey, London, ed. 2nd, 1823; https://archive.org/details/confessionsofen00dequ/page/n3/mode/2up).

3. G. F. Koob, Neurobiology of Opioid Addiction: Opponent Process, Hyperkatifeia, and Negative Reinforcement. Biol. Psychiatry 87, 44–53 (2020).

4. H. W. Matthes, R. Maldonado, F. Simonin, O. Valverde, S. Slowe, I. Kitchen, K. Befort, A. Dierich, M. Le Meur, P. Dollé, E. Tzavara, J. Hanoune, B. P. Roques, B. L. Kieffer, Loss of morphine-induced analgesia, reward effect and withdrawal symptoms in mice lacking the mu-opioid-receptor gene. Nature 383, 819–23 (1996).

5. Y. Zhu, P. Yan, R. Wang, J. Lai, H. Tang, X. Xiao, R. Yu, X. Bao, F. Zhu, K. Wang, Y. Lu, J. Dang, C. Zhu, R. Zhang, W. Dang, B. Zhang, Q. Fu, Q. Zhang, C. Kang, Y. Chen, X. Chen, Q. Liang, K. Wang, Opioid-induced fragile-like regulatory T cells contribute to withdrawal. Cell 186 (2023), doi:10.1016/j.cell.2022.12.030.

6. N. Massaly, B. A. Copits, A. R. Wilson-Poe, L. Hipólito, T. Markovic, H. J. Yoon, S. Liu, M. C. Walicki, D. L. Bhatti, S. Sirohi, A. Klaas, B. M. Walker, R. Neve, C. M. Cahill, K. I. Shoghi, R. W. Gereau, J. G. McCall, R. Al-Hasani, M. R. Bruchas, J. A. Morón, Pain-Induced Negative Affect Is Mediated via Recruitment of The Nucleus Accumbens Kappa Opioid System. Neuron 102 (2019), doi:10.1016/j.neuron.2019.02.029.

7. Y. Hayashi, S. Morinaga, J. Zhang, Y. Satoh, A. L. Meredith, T. Nakata, Z. Wu, S. Kohsaka, K. Inoue, H. Nakanishi, BK channels in microglia are required for morphine-induced hyperalgesia. Nature Communications 7 (2016), doi:10.1038/ncomms11697.

8. A.-A. Liu, P.-W. Liu, J.-W. Luo, S.-Y. Yao, B.-Y. Wang, C.-Y. Wang, R.-F. Ye, G.-Y. Zan, M. Sun, J.-G. Liu, Y.-J. Wang, Kappa opioid receptor-dependent depressive-like behaviors associated with morphine withdrawal contribute to stress-induced reinstatement of morphine seeking. Neuropsychopharmacology (2026), doi:10.1038/s41386-026-02549-1.

9. M. R. Hutchinson, Y. Shavit, P. M. Grace, K. C. Rice, S. F. Maier, L. R. Watkins, Exploring the neuroimmunopharmacology of opioids: An integrative review of mechanisms of central immune signaling and their implications for opioid analgesia. Pharmacol. Rev. 63 (2011), doi:10.1124/pr.110.004135.

10. Y.-I. Hser, L. J. Mooney, A. J. Saxon, K. Miotto, D. S. Bell, D. Huang, Chronic pain among patients with opioid use disorder: Results from electronic health records data. J. Subst. Abuse Treat. 77, 26–30 (2017).

11. R. N. Bluthenthal, K. Simpson, R. C. Ceasar, J. Zhao, L. Wenger, A. H. Kral, Opioid withdrawal symptoms, frequency, and pain characteristics as correlates of health risk among people who inject drugs. Drug Alcohol Depend. 211, 107932 (2020).

12. L. M. Rieb, K. DeBeck, K. Hayashi, E. Wood, E. Nosova, M. J. Milloy, Withdrawal-associated injury site pain prevalence and correlates among opioid-using people who inject drugs in Vancouver, Canada. Drug Alcohol Depend. 216 (2020), doi:10.1016/j.drugalcdep.2020.108242.

13. L. M. Rieb, W. V. Norman, R. E. Martin, J. Berkowitz, E. Wood, M. J. Milloy, R. McNeil, Linking opioid-induced hyperalgesia and withdrawal-associated injury site pain: A case report. Pain Rep. 3 (2018), doi:10.1097/PR9.0000000000000648.

14. P. M. Grace, M. R. Hutchinson, S. F. Maier, L. R. Watkins, Pathological pain and the neuroimmune interfaceNat. Rev. Immunol. 14 (2014), doi:10.1038/nri3621.

15. Government of Canada, Opioid- and Stimulant-related Harms in Canadahttps://health-infobase.canada.ca/substance-related-harms/opioids-stimulants/ (2026).

16. Statistics Canada, Table 13-10-0392-01 Deaths and age-specific mortality rates, by selected grouped causesdoi: 10.25318/1310039201-eng (2025).

17. D. Dowell, S. Brown, S. Gyawali, J. Hoenig, J. Ko, C. Mikosz, E. Ussery, G. Baldwin, C. M. Jones, Y. Olsen, N. Tomoyasu, B. Han, W. M. Compton, N. D. Volkow, Treatment for Opioid Use Disorder: Population Estimates — United States, 2022. MMWR Morb. Mortal. Wkly. Rep. 73 (2024), doi:10.15585/mmwr.mm7325a1.

18. S. P. Stumbo, B. J. H. Yarborough, D. McCarty, C. Weisner, C. A. Green, Patient-reported pathways to opioid use disorders and pain-related barriers to treatment engagement. J. Subst. Abuse Treat. 73, 47–54 (2017).

19. I. Sankaranarayanan, K. Mazhar, A. M. Barry, S. Shiers, A. Wangzhou, J. R. Sondermann, F. Xian, N. Inturi, M. A. Wilde, E. C. Meyers, J. B. Lesnak, J. A. O’Brien, M. S. Yousuf, M. Schmidt, G. Dussor, E. Vines, P. Horton, A. Cervantes, T. Khan, G. Funk, D. Tavares-Ferreira, PRECISION Human Pain Network, P. M. Dougherty, M. Curatolo, T. J. Price, Progressive neurodegeneration in human dorsal root ganglion from diabetes to painful neuropathy. bioRxiv (2026), doi:10.64898/2026.01.16.700028.

20. A. M. Barry, J. R. Sondermann, J. B. Lesnak, F. Xian, Ú. Franco-Enzástiga, J. A. O’Brien, D. Gomez-Varela, M. K. Schackmuth, S. Shiers, T. J. Price, M. Schmidt, Multi-omic integration with human dorsal root ganglia proteomics highlights TNFα signalling as a relevant sexually dimorphic pathway. Pain 166 (2025), doi:10.1097/j.pain.0000000000003656.

21. E. S. Staedtler, M. R. Sapio, D. M. King, D. Maric, A. Ghetti, A. J. Mannes, M. J. Iadarola, The μ-opioid receptor differentiates two distinct human nociceptive populations relevant to clinical pain. Cell Rep. Med. 5, 101788 (2024).

22. L. R. Watkins, M. R. Hutchinson, K. C. Rice, S. F. Maier, The “Toll” of Opioid-Induced Glial Activation: Improving the Clinical Efficacy of Opioids by Targeting GliaTrends Pharmacol. Sci. 30 (2009), doi:10.1016/j.tips.2009.08.002.

23. S. A. Bhuiyan, S. S. Nagi, I. Sankaranarayanan, E. Semizoglou, D. Usoskin, L. Yang, H. Yu, A. Arendt-Tranholm, Z. Bertels, P. Bhatia, O. Bouchatta, K. Boyer, A. Cervantes, J. Chalif, H. Chintalapudi, A. Cicalo, B. Copits, C. Cronin, M. Curatolo, X. Dong, P. M. Dougherty, A. Dourson, G. Funk, K. Gabriel, D. S. Griesemer, H. Guo, P. Gupta, C. Hofstetter, P. Horton, A. Hsieh, N. N. Inturi, A. Jain, S. Jayakar, B. Johnston, R. Kim, D. Krauter, J. Kupari, J. Lemen, J. B. Lesnak, W. Liu, I. Lopez, Y. Lu, H. J. MacMillan, K. Mazhar, P. Meriau, J. R. Moffitt, M. M. Moreno, J. M. Mwirigi, H. Naz, J. O’Brein, M. Payne, J. Del Rosario, S. F. Rosen, S. Shiers, E. Simpson, R. Slivicki, J. R. Stone, D. Tavares-Ferreira, M. Uhelski, C. J. Woolf, Q. Xu, J. Yi, M. S. Yousuf, D. Zhu, V. Cavalli, G. Zhao, H. Olausson, P. Ernfors, R. W. Gereau, W. Luo, T. J. Price, W. Renthal, NIH PRECISION Human Pain Network, A Reference Atlas of the Human Dorsal Root Ganglion. bioRxiv (2025), doi:10.1101/2025.11.05.686654.

24. M. I. Love, W. Huber, S. Anders, Moderated estimation of fold change and dispersion for RNA-seq data with DESeq2. Genome Biol. 15 (2014), doi:10.1186/s13059-014-0550-8.

25. A. Zhu, J. G. Ibrahim, M. I. Love, Heavy-Tailed prior distributions for sequence count data: Removing the noise and preserving large differences. Bioinformatics 35 (2019), doi:10.1093/bioinformatics/bty895.

26. S. F. Wolf, P. A. Temple, M. Kobayashi, D. Young, M. Dicig, L. Lowe, R. Dzialo, L. Fitz, C. Ferenz, R. M. Hewick, Cloning of cDNA for natural killer cell stimulatory factor, a heterodimeric cytokine with multiple biologic effects on T and natural killer cells. The Journal of Immunology 146 (1991), doi:10.4049/jimmunol.146.9.3074.

27. K. Pekayvaz, C. Gold, P. Hoseinpour, A. Engel, A. Martinez-Navarro, L. Eivers, R. Coletti, M. Joppich, F. Dionísio, R. Kaiser, L. Tomas, A. Janjic, M. Knott, F. Mehari, V. Polewka, M. Kirschner, A. Boda, L. Nicolai, H. Schulz, A. Titova, B. Kilani, M. Lorenz, G. Fingerle-Rowson, R. Bucala, W. Enard, R. Zimmer, C. Weber, P. Libby, C. Schulz, S. Massberg, K. Stark, Mural cell-derived chemokines provide a protective niche to safeguard vascular macrophages and limit chronic inflammation. Immunity 56 (2023), doi:10.1016/j.immuni.2023.08.002.

28. A. Büttner, K. Rohrmoser, G. Mall, R. Penning, S. Weis, Widespread axonal damage in the brain of drug abusers as evidenced by accumulation of β-amyloid precursor protein (β-APP): An immunohistochemical investigation. Addiction 101 (2006), doi:10.1111/j.1360-0443.2006.01505.x.

29. I. C. Anthony, K. E. Norrby, T. Dingwall, F. W. Carnie, T. Millar, J. C. Arango, R. Robertson, J. E. Bell, Predisposition to accelerated Alzheimer-related changes in the brains of human immunodeficiency virus negative opiate abusers. Brain 133 (2010), doi:10.1093/brain/awq263.

30. S. N. Ramage, I. C. Anthony, F. W. Carnie, A. Busuttil, R. Robertson, J. E. Bell, Hyperphosphorylated tau and amyloid precursor protein deposition is increased in the brains of young drug abusers. Neuropathol. Appl. Neurobiol. 31 (2005), doi:10.1111/j.1365-2990.2005.00670.x.

31. D. Saul, R. L. Kosinsky, E. J. Atkinson, M. L. Doolittle, X. Zhang, N. K. LeBrasseur, R. J. Pignolo, P. D. Robbins, L. J. Niedernhofer, Y. Ikeno, D. Jurk, J. F. Passos, L. T. J. Hickson, A. Xue, D. G. Monroe, T. Tchkonia, J. L. Kirkland, J. N. Farr, S. Khosla, A new gene set identifies senescent cells and predicts senescence-associated pathways across tissues. Nature Communications 13 (2022), doi:10.1038/s41467-022-32552-1.

32. L. M. Snyder, M. C. Chiang, E. Loeza-Alcocer, Y. Omori, J. Hachisuka, T. D. Sheahan, J. R. Gale, P. C. Adelman, E. I. Sypek, S. A. Fulton, R. L. Friedman, M. C. Wright, M. G. Duque, Y. S. Lee, Z. Hu, H. Huang, X. Cai, K. A. Meerschaert, V. Nagarajan, T. Hirai, G. Scherrer, D. H. Kaplan, F. Porreca, B. M. Davis, M. S. Gold, H. R. Koerber, S. E. Ross, Kappa Opioid Receptor Distribution and Function in Primary Afferents. Neuron 99 (2018), doi:10.1016/j.neuron.2018.08.044.

33. S. A. Bhuiyan, M. Xu, L. Yang, E. Semizoglou, P. Bhatia, K. I. Pantaleo, I. Tochitsky, A. Jain, B. Erdogan, S. Blair, V. Cat, J. M. Mwirigi, I. Sankaranarayanan, D. Tavares-Ferreira, U. Green, L. A. McIlvried, B. A. Copits, Z. Bertels, J. S. Del Rosario, A. J. Widman, R. A. Slivicki, J. Yi, R. Sharif-Naeini, C. J. Woolf, J. K. Lennerz, J. L. Whited, T. J. Price, R. W. Gereau, W. Renthal, Harmonized cross-species cell atlases of trigeminal and dorsal root ganglia. Sci. Adv. 10 (2024), doi:10.1126/sciadv.adj9173.

34. Ú. Franco-Enzástiga, K. Natarajan, F. Espinosa, R. Granja-Vazquez, H. Mydugolam, T. J. Price, Type I IFNs enhance human dorsal root ganglion nociceptor excitability and induce TRPV1 sensitization. JCI Insight 10 (2025), doi:10.1172/jci.insight.194987.

35. L. C. Colodeti, T. B. P. Gonçalves, H. P. Mota-Araujo, S. M. B. Araujo, G. N. Pires, B. V. M. Santiago, T. S. Bacelar, F. L. Fontes-Dantas, E. C. C. Rennô, M. O. L. da Silva, L. E. B. Savio, P. S. Ciriaco, T. B. Taboada, A. B. O. Barbosa, F. M. Gomes, A. M. B. Martinez, L. J. Costa, I. Assunção-Miranda, N. R. Villela, J. L. Proença-Modena, H. R. Mendonça, S. Alves-Leon, G. F. Passos, W. M. de Souza, R. da Costa, C. P. Figueiredo, Acute IFN-γ responses drive sensory neuron injury and chronic pain after chikungunya virus infection. bioRxiv (2026), doi:10.64898/2026.02.02.702518.

36. J. Serra, H. Bostock, R. Solà, J. Aleu, E. García, B. Cokic, X. Navarro, C. Quiles, Microneurographic identification of spontaneous activity in C-nociceptors in neuropathic pain states in humans and rats. Pain 153 (2012), doi:10.1016/j.pain.2011.08.015.

37. R. Y. North, Y. Li, P. Ray, L. D. Rhines, C. E. Tatsui, G. Rao, C. A. Johansson, H. Zhang, Y. H. Kim, B. Zhang, G. Dussor, T. H. Kim, T. J. Price, P. M. Dougherty, Electrophysiological and transcriptomic correlates of neuropathic pain in human dorsal root ganglion neurons. Brain 142 (2019), doi:10.1093/brain/awz063.

38. F. Hawkins, in (Burgess and Hill, London, 1826), pp. 5–347.

39. M. L. Uhelski, M. K. Schaub, F. Espinosa, M. Heles, N. Cortes, Y. Li, C. E. Tatsui, L. D. Rhines, R. Y. North, C. Alvarez-Breckenridge, G. Funk, P. Horton, A. Cervantes, J. B. Lesnak, M. Curatolo, G. Dussor, T. J. Price, P. M. Dougherty, Suzetrigine (VX-548) exhibits activity-dependent effects on human dorsal root ganglion neurons with differential pain etiology. Pain 167, 1430–1439 (2026).

40. M. L. Seney, S. M. Kim, J. R. Glausier, M. A. Hildebrand, X. Xue, W. Zong, J. Wang, M. A. Shelton, B. D. N. Phan, C. Srinivasan, A. R. Pfenning, G. C. Tseng, D. A. Lewis, Z. Freyberg, R. W. Logan, Transcriptional Alterations in Dorsolateral Prefrontal Cortex and Nucleus Accumbens Implicate Neuroinflammation and Synaptic Remodeling in Opioid Use Disorder. Biol. Psychiatry 90 (2021), doi:10.1016/j.biopsych.2021.06.007.

41. G. L. Riß, D. I. Chang, C. Wevers, A. M. Westendorf, J. Buer, N. Scherbaum, W. Hansen, Opioid maintenance therapy restores CD4+ T cell function by normalizing CD4+CD25high regulatory T cell frequencies in heroin user. Brain Behav. Immun. 26 (2012), doi:10.1016/j.bbi.2012.05.008.

42. I. N. Johnston, E. D. Milligan, J. Wieseler-Frank, M. G. Frank, V. Zapata, J. Campisi, S. Langer, D. Martin, P. Green, M. Fleshner, L. Leinwand, S. F. Maier, L. R. Watkins, A role for proinflammatory cytokines and fractalkine in analgesia, tolerance, and subsequent pain facilitation induced by chronic intrathecal morphine. Journal of Neuroscience 24 (2004), doi:10.1523/JNEUROSCI.1850-04.2004.

43. Y. Cui, X. X. Liao, W. Liu, R. X. Guo, Z. Z. Wu, C. M. Zhao, P. X. Chen, J. Q. Feng, A novel role of minocycline: Attenuating morphine antinociceptive tolerance by inhibition of p38 MAPK in the activated spinal microglia. Brain Behav. Immun. 22 (2008), doi:10.1016/j.bbi.2007.07.014.

44. P. Song, Z. Q. Zhao, The involvement of glial cells in the development of morphine tolerance. Neurosci. Res. 39 (2001), doi:10.1016/S0168-0102(00)00226-1.

45. P. Nagyoszi, I. Wilhelm, A. E. Farkas, C. Fazakas, N. T. K. Dung, J. Haskó, I. A. Krizbai, Expression and regulation of toll-like receptors in cerebral endothelial cells. Neurochem. Int. 57 (2010), doi:10.1016/j.neuint.2010.07.002.

46. A. H. Klein, S. M. S. Alam, K. Johnson, C. Kriner, B. Beck, B. Nelson, C. Hill, B. Meyer, J. Mellang, V. J. Watts, Inhibition of adenylyl cyclase 1 or exchange protein activated by cAMP restores ATP-sensitive potassium channel activity after chronic opioid exposure. Br. J. Pharmacol. 183 (2026), doi:10.1111/bph.70417.

47. M. Iftinca, L. Basso, R. Flynn, C. Kwok, C. Roland, A. Hassan, M. Defaye, R. Ramachandran, T. Trang, C. Altier, Chronic morphine regulates TRPM8 channels via MOR-PKCβ signaling. Mol. Brain 13 (2020), doi:10.1186/s13041-020-00599-0.

48. D. Araldi, O. Bogen, P. G. Green, J. D. Levine, Role of nociceptor toll-like receptor 4 (tlr4) in opioid-induced hyperalgesia and hyperalgesic priming. Journal of Neuroscience 39 (2019), doi:10.1523/JNEUROSCI.0966-19.2019.

49. E. V. Khomula, D. Araldi, P. G. Green, J. D. Levine, Sensitization of human and rat nociceptors by low dose morphine is toll-like receptor 4-dependent. Mol. Pain 20 (2024), doi:10.1177/17448069241227922.

50. W. Ma, W. H. Zheng, S. Kar, R. Quirion, Morphine treatment induced calcitonin gene-related peptide and substance P increases in cultured dorsal root ganglion neurons. Neuroscience 99 (2000), doi:10.1016/S0306-4522(00)00226-8.

51. N. M. Wilson, H. Jung, M. S. Ripsch, R. J. Miller, F. A. White, CXCR4 signaling mediates morphine-induced tactile hyperalgesia. Brain Behav. Immun. 25 (2011), doi:10.1016/j.bbi.2010.12.014.

52. S. Yamakita, D. Fujita, K. Sudo, D. Ishikawa, K. Kushimoto, Y. Horii, F. Amaya, Activation of neurons and satellite glial cells in the DRG produces morphine-induced hyperalgesia. Mol. Pain 19 (2023), doi:10.1177/17448069231181973.

53. J. Su, M. D. Zhang, J. Kupari, D. Kwak, L. Picton, B. Xu, L. F. do Nascimento, Y. Hu, A. Gonzalez, D. Usoskin, Z. Xu, M. Szczot, A. El Manira, R. Holmdahl, P. Ernfors, Persistent interferon signaling causes sensory neuron plasticity and pain before and during arthritis. Nat. Neurosci. (2026), doi:10.1038/s41593-026-02234-y.

54. L. R. Watkins, S. F. Maier, L. E. Goehler, Immune activation: the role of pro-inflammatory cytokines in inflammation, illness responses and pathological pain statesPain 63 (1995), doi:10.1016/0304-3959(95)00186-7.

55. T. Hucho, J. D. Levine, Signaling Pathways in Sensitization: Toward a Nociceptor Cell BiologyNeuron 55 (2007), doi:10.1016/j.neuron.2007.07.008.

56. M. Trøstheim, M. Eikemo, Hyperalgesia in Patients with a History of Opioid Use Disorder: A Systematic Review and Meta-Analysis. JAMA Psychiatry 81 (2024), doi:10.1001/jamapsychiatry.2024.2176.

57. K. D. Ersche, L. Clark, M. London, T. W. Robbins, B. J. Sahakian, Profile of executive and memory function associated with amphetamine and opiate dependence. Neuropsychopharmacology 31 (2006), doi:10.1038/sj.npp.1300889.

58. J. J. Michel, K. Sanghvi, J. Rosenbauer, L. Humbs, C. T. Dierssen, S. Grudzenski-Theis, V. Sachs, K. Jähne, K. Degenhardt, L. Frölich, J. Herms, M. Fatar, M. Platten, L. Bunse, Type I interferon drives T cell responses to amyloid beta in the central nervous system. Nature Communications 17 (2026), doi:10.1038/s41467-026-72262-6.

59. K. K. Ting, P. Coleman, H. J. Kim, Y. Zhao, J. Mulangala, N. C. Cheng, W. Li, D. Gunatilake, D. M. Johnstone, L. Loo, G. G. Neely, P. Yang, J. Götz, M. A. Vadas, J. R. Gamble, Vascular senescence and leak are features of the early breakdown of the blood–brain barrier in Alzheimer’s disease models. Geroscience 45 (2023), doi:10.1007/s11357-023-00927-x.

60. N. D. Volkow, C. Blanco, Substance use disorders: a comprehensive update of classification, epidemiology, neurobiology, clinical aspects, treatment and prevention. World Psychiatry 22 (2023), doi:10.1002/wps.21073.

61. E. R. Butelman, Y. Huang, F. Cathomas, P. O. Gaudreault, P. Roussos, S. J. Russo, R. Z. Goldstein, N. Alia-Klein, Serum cytokines and inflammatory proteins in individuals with heroin use disorder: potential mechanistically based biomarkers for diagnosis. Transl. Psychiatry 14 (2024), doi:10.1038/s41398-024-03119-z.

62. S. Shiers, M. Saad Yousuf, J. Mwirigi, A. Cervantes, T. Price, Human Ganglia and Spinal Cord Tissue Procurement from Organ Donors and Tissue Quality Assessment v1 (2024), doi:10.17504/protocols.io.kqdg32qr1v25/v1.

63. J. B. Lesnak, M. K. Schaub, K. Gomez, A. Calderon-Rivera, S. Loya-Lopez, R. Stewart, S. Jo, A. Fujita, T. Osorno, H. Mydugolam, M. Desai, K. Natarajan, M. K. Schackmuth, M. M. Moreno, S. I. Shiers, A. Cervantes, G. Funk, P. Horton, E. Vines, M. S. Yousuf, K. E. Sadler, B. P. Bean, R. Khanna, G. Dussor, T. J. Price, An improved preservation method for human dorsal root ganglion neurons enables wider access to human molecular pain neuroscience. Cell reports methods 6, 101412 (2026).

64. M. V. Valtcheva, B. A. Copits, S. Davidson, T. D. Sheahan, M. Y. Pullen, J. G. McCall, K. Dikranian, R. W. Gereau, Surgical extraction of human dorsal root ganglia from organ donors and preparation of primary sensory neuron cultures. Nat. Protoc. 11 (2016), doi:10.1038/nprot.2016.111.

65. N. P. du Sert, A. Ahluwalia, S. Alam, M. T. Avey, M. Baker, W. J. Browne, A. Clark, I. C. Cuthill, U. Dirnagl, M. Emerson, P. Garner, S. T. Holgate, D. W. Howells, V. Hurst, N. A. Karp, S. E. Lazic, K. Lidster, C. J. MacCallum, M. Macleod, E. J. Pearl, O. H. Petersen, F. Rawle, P. Reynolds, K. Rooney, E. S. Sena, S. D. Silberberg, T. Steckler, H. Würbel, Reporting animal research: Explanation and elaboration for the arrive guidelines 2.0PLoS Biol. 18 (2020), doi:10.1371/journal.pbio.3000411.

66. A. Silveira Prudente, S. Hoon Lee, J. Roh, D. D. Luckemeyer, C. F. Cohen, M. Pertin, C. K. Park, M. R. Suter, I. Decosterd, J. M. Zhang, R. R. Ji, T. Berta, Microglial STING activation alleviates nerve injury-induced neuropathic pain in male but not female mice. Brain Behav. Immun. 117 (2024), doi:10.1016/j.bbi.2024.01.003.

67. A. Barry, T. Price, M. Schmidt, Bulk Proteomics (DIA-MS) of Human Dorsal Root Ganglion v1 (2024), doi:10.17504/protocols.io.j8nlk8j56l5r/v1.

68. A. Barry, T. Price, M. Schmidt, Bulk Proteomics (DIA-MS) of Human Dorsal Root Ganglion v1 (2024), doi:10.17504/protocols.io.j8nlk8j56l5r/v1.

69. V. Demichev, C. B. Messner, S. I. Vernardis, K. S. Lilley, M. Ralser, DIA-NN: neural networks and interference correction enable deep proteome coverage in high throughput. Nat. Methods 17 (2020), doi:10.1038/s41592-019-0638-x.

70. M. E. Ritchie, B. Phipson, D. Wu, Y. Hu, C. W. Law, W. Shi, G. K. Smyth, Limma powers differential expression analyses for RNA-sequencing and microarray studies. Nucleic Acids Res. 43 (2015), doi:10.1093/nar/gkv007.

71. D. Wu, G. K. Smyth, Camera: A competitive gene set test accounting for inter-gene correlation. Nucleic Acids Res. 40 (2012), doi:10.1093/nar/gks461.

72. T. Wu, E. Hu, S. Xu, M. Chen, P. Guo, Z. Dai, T. Feng, L. Zhou, W. Tang, L. Zhan, X. Fu, S. Liu, X. Bo, G. Yu, clusterProfiler 4.0: A universal enrichment tool for interpreting omics data. Innovation 2 (2021), doi:10.1016/j.xinn.2021.100141.

73. P. L. Germain, M. D. Robinson, A. Lun, C. Garcia Meixide, W. Macnair, Doublet identification in single-cell sequencing data using scDblFinder. F1000Res. 10 (2022), doi:10.12688/f1000research.73600.2.

74. M. D. Young, S. Behjati, SoupX removes ambient RNA contamination from droplet-based single-cell RNA sequencing data. Gigascience 9 (2020), doi:10.1093/gigascience/giaa151.

75. S. Jin, C. F. Guerrero-Juarez, L. Zhang, I. Chang, R. Ramos, C. H. Kuan, P. Myung, M. V. Plikus, Q. Nie, Inference and analysis of cell-cell communication using CellChat. Nat. Commun. 12 (2021), doi:10.1038/s41467-021-21246-9.

76. J. W. Squair, M. Gautier, C. Kathe, M. A. Anderson, N. D. James, T. H. Hutson, R. Hudelle, T. Qaiser, K. J. E. Matson, Q. Barraud, A. J. Levine, G. La Manno, M. A. Skinnider, G. Courtine, Confronting false discoveries in single-cell differential expression. Nat. Commun. 12, 5692 (2021).

77. K. M. Cahill, Z. Huo, G. C. Tseng, R. W. Logan, M. L. Seney, Improved identification of concordant and discordant gene expression signatures using an updated rank-rank hypergeometric overlap approach. Sci. Rep. 8 (2018), doi:10.1038/s41598-018-27903-2.

78. Y. Perez-Riverol, C. Bandla, D. J. Kundu, S. Kamatchinathan, J. Bai, S. Hewapathirana, N. S. John, A. Prakash, M. Walzer, S. Wang, J. A. Vizcaíno, The PRIDE database at 20 years: 2025 update. Nucleic Acids Res. 53 (2025), doi:10.1093/nar/gkae1011.

